# Dietary bacteria control *C. elegans* fat content through pathways converging at phosphatidylcholine

**DOI:** 10.1101/2024.02.24.581900

**Authors:** Hsiao-Fen Han, Shao-Fu Nien, Hang-Shiang Jiang, Jui-Ching Wu, Chia-Yi Chiang, Man-Tzu Li, Leng-Jie Huang, Sufeng Chiang, Lien-Chieh Lin, Yi-Ting Chuang, Yu-Ho Lin, Chao-Wen Wang, Yi-Chun Wu

## Abstract

Dietary factors play a pivotal role in regulating metabolism in both health and disease. Lipid metabolism is particularly important for organismal health and longevity. However, the mechanisms by which dietary factors influence lipid metabolism remain poorly understood. Here, using the nematode *C. elegans* as a model system, we investigated the influence of distinct bacterial diets on fat metabolism. We found that dietary vitamin B12 activates the S-adenosyl methionine (SAM) and phosphatidylcholine (PC) biosynthetic pathways. This activation leads to elevated levels of PC, which in turn suppresses the expression of the gene *fat-7* and modulates lipid droplet dynamics through the regulatory proteins SBP-1/SREBP1 and SEIP-1/SEIPIN, respectively. Additionally, we identified a feedback loop involving SBP-1-mediated regulation of acid sphingomyelinase ASM-3, which enhances the production of phospho-choline and further stimulates PC synthesis. Our localization studies further suggest that ASM-3 may act as a signaling mediator between the intestine and coelomocytes, coordinating their roles in vitamin B12-mediated fat regulation. Overall, our findings shed new light on the complex interplay between diet and metabolic regulation, with a particular emphasis on the central role of phosphatidylcholine.

**Highlights:**

- Animals govern PC level to regulate lipid homeostasis in response to diets
- B12 regulates SAM-PC axis to affect lipogenic genes expression and LD biogenesis
- Coelomocytes regulate diets-induced lipid homeostasis through *asm-3*
- *asm-3* constructs a positive feedback loop to participate in PC metabolism

## Introduction

Dietary factors are increasingly recognized as key players in the onset and prevention of major diseases, such as cancers, coronary heart diseases, and diabetes^1,2^. Recent research indicates that diet not only shapes nutrient intake but also influences the intestinal microbiome, and microbiome-derived metabolites, in turn, impact organismal metabolism and immune responses^3^. However, the direct or indirect mechanisms through which dietary composition influences host gene expression, biological processes, and physiological states in most cases remain unclear. The nematode *C. elegans* serves as an excellent model for dietary studies due to its well-characterized genetics^4^, conserved metabolic pathways^5,6^, and simplified feeding regimen consisting of a bacterial monoculture^7^. Recent work in *C. elegans* has uncovered several mechanisms by which diet modulates important physiological processes, such as growth rate and longevity^8–16^.

Vitamin B12 is synthesized by some bacteria and archaea, but not by plants or animals^17^. Thus, B12 is transferred to and accumulated inside the body through dietary sources or microbial interaction^17^. B12 exerts its influence primarily through two distinct pathways to regulate physiology^16,18^. Pathway I regulates the one-carbon (1C) cycle that promotes the synthesis of the methyl donor group S-adenosyl methionine (SAM). Through the methylation of DNA, histones and other proteins, SAM levels influence many physiological processes including growth rate, and fertility^19,20^. Pathway II promotes the breakdown of toxic propionyl-CoA, an intermediate in the catabolism of odd-number fatty acids and branched-chain amino acids^11,16,21^. Through these metabolic effects, B12 promotes faster development and affects lifespan^11,16,18,21,22^.

An intricate system of lipogenesis, lipolysis, lipophagy, lipid transport, and lipid utilization contributes to lipid homeostasis, which is crucial for a wide range of physiological functions such as energy storage, membrane structure, signaling transduction, and lipid-based hormone production^23–25^. Apart from direct dietary lipid intake, the mechanistic impact of other diet components on organismal lipid metabolism remains poorly understood.

In this study, we investigated the mechanisms underlying dietary control of host lipid metabolism, using *C. elegans* fed one of two distinct bacteria diets. Lipidomic and transcriptomic analyses pointed to a pivotal role for *fat-7,* a gene encoding a Δ9 fatty acid desaturase, in the striking reduction in organismal lipid content we observed when worms were fed a non-canonical bacterial diet. Mechanistically, we show that dietary vitamin B12, enriched in one bacterial diet but not the other, stimulates SAM production and phosphatidylcholine (PC) synthesis. In turn, the SAM-PC axis negatively regulates *fat-7* in a SBP-1-dependent manner, leading to suppressed lipogenesis in response to dietary vitamin B12. Elevated PC levels impact intestinal lipid droplet (LD) dynamics by hindering the recruitment of SEIP-1 to peri-LD cages. We also show that signaling orchestrated by the acid sphingomyelinase, ASM-3 converges at the PC synthesis pathway to further bolster B12-mediated lipid reduction. Altogether, our data elucidate how the micronutrient B12 from bacteria utilizes PC as the convergence point for multiple regulatory pathways to tune lipid homeostasis in response to diets, providing valuable insights for dietary interventions in managing lipid related disorders.

## Results

### Diet affects host lipid level

To study how host lipid homeostasis is influenced by diets, we measured the lipid content of *C. elegans* fed two different bacterial diets – *Comamonas aquatica* DA1877 (hereafter referred to as DA) or the standard *Escherichia coli* OP50 (hereafter, OP) - using Oil Red O (O.R.O) staining and thin layer chromatography (TLC). We found that DA-fed worm had a significantly decreased level of neutral lipids, including triacylglycerol (TAG), in comparison to OP-fed worms (Figure 1A, B).

**Figure 1.**
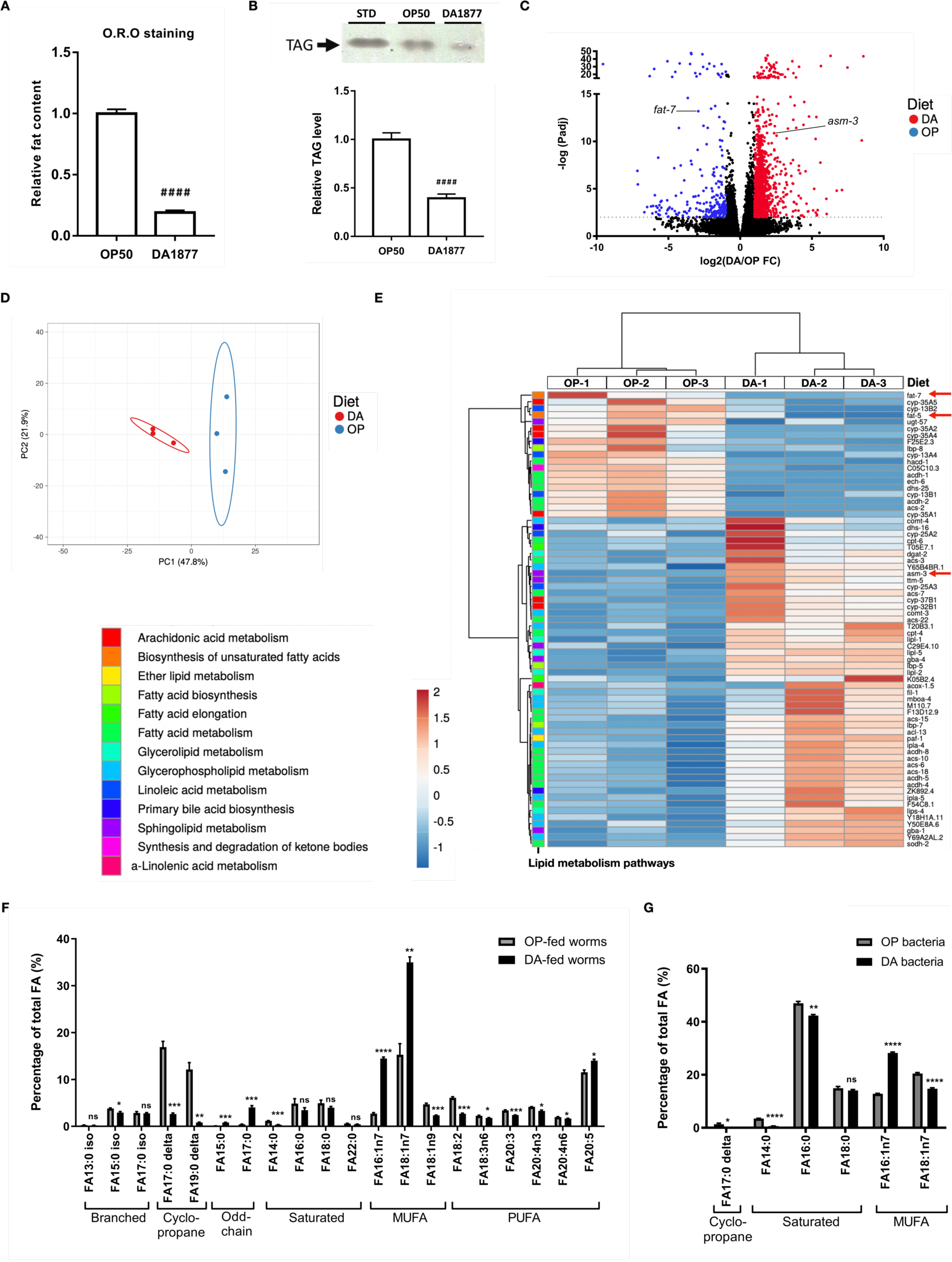
DA1877 diet represses lipid content and induces lipid metabolic reprogramming in *C. elegans*. (A) O.R.O staining results of wild-type worms on OP or DA diet. (B) TLC analysis of TAG level in wild-type animals feeding OP or DA. (C) Volcano plot of RNA-seq results. The expression is calculated based on TPM (transcripts per million) and each dot represents one gene. Black dots, red dots and blue dots represent genes with similar expression level, upregulated level, and downregulated level in DA-fed worms compared to OP-fed animals, respectively. (D) Principal component analysis of the expression of 471 lipid metabolic genes in OP- and DA-fed worms. Gene expression data was from RNA-seq analysis. (E) Heatmap for the differentially expressed metabolic genes involved in 13 lipid metabolism pathways between OP- and DA-fed worms. The graph was generated using ClustVis (https://biit.cs.ut.ee/clustvis/). (F) Fatty acid composition of OP- and DA-fed worms by GC-MS. The fatty acid content is shown as percentage of total fatty acids. (G) GC-MS analysis of fatty acids in OP and DA bacteria. Statistical analyses in (A), (B), (F) & (G) were performed using Two-tailed T tests and the data is presented as mean ± SEM. # indicates comparison of dietary effects between OP- and DA-fed wild-type worms. * indicates comparison of individual fatty acid percentage between different diets-fed worms or different bacteria. * ^/#^ *P* < 0.05, ** ^/##^ *P* < 0.01, *** ^/###^ *P* < 0.001, **** ^/####^ *P* < 0.0001.

Decreased food intake is known to reduce organismal lipid content^26^. To examine whether the altered lipid level we observed in DA-fed worms is caused by a discrepancy in food intake, we measured the pharyngeal pumping rate of worms fed each of the bacterial diets and found them to be very similar (Supplementary Figure S1A). Given that *C. elegans* gustatory and olfactory perceptions have been shown to play active roles in regulating lipid content^27,28,29^, we also tested whether the reduced lipid content is mediated by the sensing of DA by amphid neurons. The lipid content in *daf-6* mutants - in which the formation of amphid sheath cells is abnormal^30^ - is comparable with wild-type worms across both diet conditions (Supplementary Figure S1B), suggesting that neuronal sensing of bacteria was not responsible for the observed effect.

To understand the mechanism underlying the regulation of host lipid content triggered by DA, we examined the gene expression changes elicited by the two different bacterial diets in young adult animals by RNA-seq. Strikingly, we found 1811 differentially expressed genes (DEGs; ≥2 or ≤-2 fold; *Padj* < 0.01) between worms fed the two different bacterial diets (Figure 1C). KEGG pathway analysis revealed the DEGs are enriched in fatty acid metabolism (*P value* = 8.84E-8). We examined a comprehensive list of 471 lipid metabolic genes^31^ and found by principal component analysis (PCA) that these genes exhibited clear separation in DA- and OP-fed worms (Figure 1D). Notably, 13 out of the 16 lipid metabolic pathways are affected by DA diet (Figure 1E). In particular, genes related to the biosynthesis of unsaturated fatty acids showed a significant decrease in expression in DA-fed worms. For example, the delta-(9) fatty acid desaturases , *fat-5* and *fat-7*, (which convert fatty acids 16:0 to 16:1n7 and 18:0 to 18:1n9, respectively^32^) decreased to 33.6% and 13.3% of their original expression in DA-fed animals, respectively (Figure 1C, E).

Next, we assessed the fatty acid composition of worms fed the two diets using gas chromatography-mass spectrometry (GC-MS). Intriguingly, the amount of total fatty acids is not altered between the OP and DA diets-fed worms (Supplementary Figure S1C); rather, diet appears to substantially influence the relative levels of different fatty acid species. We found that DA results in a lower abundance of cyclopropane fatty acids (17:0 delta and 19:0 delta) and higher levels of odd- chain fatty acids (15:0 and 17:0) in the worms (Figure 1F). While the major saturated fatty acids, palmitic acid (16:0) and stearic acid (18:0), are at a similar level between DA- and OP- fed worms, the proportion of monounsaturated fatty acids (MUFAs) differs strikingly: accounting for 51.8% of the total fatty acid in DA-fed worms *vs.* 22.7% in OP-fed animals. Specifically, the level of monounsaturated n7 fatty acids, including 16:1n7 and 18:1n7, increase significantly, whereas the 18:1n9 species decreases in response to DA. Because *C. elegans* produced PUFAs from 18:1n9, as expected we also found that most polyunsaturated fatty acids (PUFAs) (18:2, 18:3, 20:3, 20:4) are at the lower level in DA-fed animals compared to OP ones (Figure 1F).

To determine whether the distinct fatty acid patterns are caused by direct ingestion of dietary lipids, we examined the fatty acid composition of DA and OP bacteria. The results indicated that the major fatty acid species present in both bacteria is 16:0 which comprises an average of 43.2% of the total fatty acid in DA, lower than its 47.0% in OP (Figure 1G). In addition, DA had undetectable level of cyclopropane fatty acid, while it was enriched in 16:1n7 (Figure 1G). Since cyclopropane fatty acids are only produced in bacteria^33^, it is likely that the low level of these compounds in DA-fed worms are attributable to their absence in the dietary bacteria. Likewise, it is plausible that the plentiful 16:1n7 in DA worms was obtained from the diet and further elongated to 18:1n7 (Figure 1F)^34^. In contrast, both DA and OP bacteria have undetectable levels of oleic acid (18:1n9) (Figure 1G), pointing to altered worm lipid metabolic pathways as the likely explanation for the observed decrease in 18:1n9 in response to DA diet. Taken together, while there are some specific instances in which dietary lipid content might explain certain observed host lipid metabolism effects, it appears that the majority of the observed changes are due to other, indirect effect of diet on organismal physiology.

### Genetic screen for modulators of diet-mediated lipid content

Our combined transcriptomic and lipidomic analyses revealed that worms have significantly reduced expression of *fat-7* and its product, oleic acid (18:1n9) in response to the DA diet (Figure 1C, E, F), which we validated by qRT-PCR (Figure 2A) and in worms expressing a FAT-7::GFP translational reporter (Figure 2B). Overexpression of *fat-7* in the intestine of DA-fed worms increased their lipid content, while *fat-7(wa36)* mutant animals fed OP bacteria exhibited reduced lipid content (Figure 2C). These data demonstrate the sufficiency and necessity of *fat-7* for the observed diet-mediated changes in lipid content.

**Figure 2.**
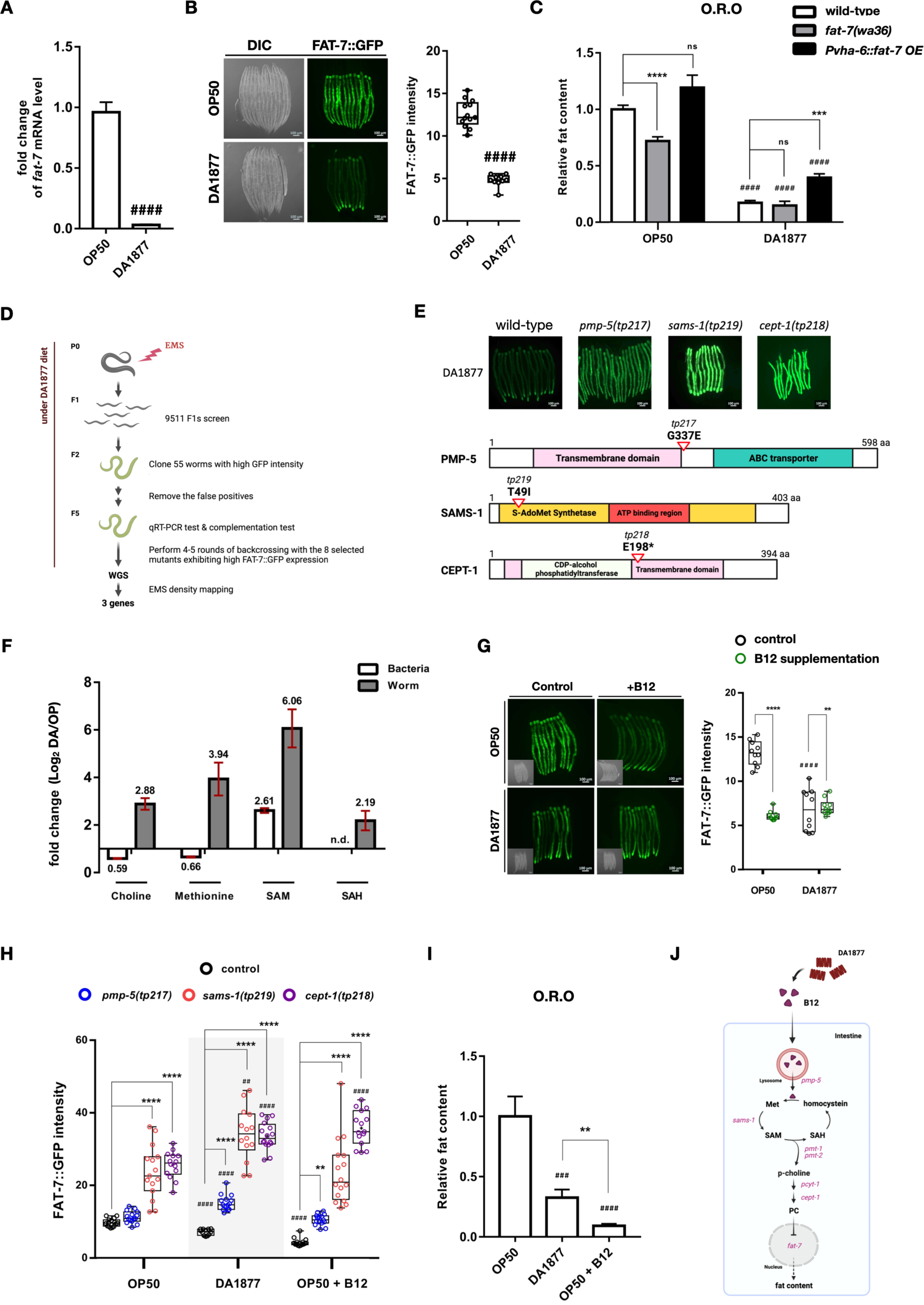
DA diet decreases *fat-7* expression level and lipid content through B12-SAM-PC axis. (A) qRT-PCR results of *fat-7* mRNA level in the worms feeding OP or DA. Experiment contained three biological repeats and n = 10 for each repeat. (B) Representative images and quantitative results of FAT-7::GFP in OP or DA-fed worms. n ≥ 10 for each sample. (C) O.R.O staining results of wild-type, *fat-7(wa36)* mutants and *tpEx697* worms, where *fat-7* is overexpressed by using intestine-specific promotor *vha-6* under OP or DA feeding. Experiment contained 2-3 biological repeats and n ≥ 50 in each repeat. (D) Flowchart of forward genetic screen. (E) Upper panel images showing FAT-7::GFP expression of wild-type animals and the isolated mutants from forward genetic screen on DA diet. The lower scheme illustrating the amino acid changes in those identified alleles. (F) LC/MS/MS analysis of metabolites extracted from worms (gray bar) and bacteria (white bar). n.d. means undetectable. (G) Images and quantitative results of FAT-7::GFP signals in FAT-7 reporter strain DMS303(*nIs590*) feeding OP or DA with or without B12 supplementation. (H) Quantitative results of FAT-7::GFP signals in the control worms DMS303(*nIs590)* and mutants isolated from screen feeding OP, DA, or OP with B12 supplementation. (I) O.R.O staining results of wild-type worms feeding OP, DA or OP supplemented with 64nM B12. (J) Cartoon showing the mechanism of B12 in regulation of fat content under DA diet. The graphic work in (D), (E), (J) are created with BioRender.com. Statistical analysis was performed using Two-tailed T tests and the bar plot is presented as mean ± SEM. The centerline in the box of boxplot denotes the median value in the data. # indicates comparison of dietary effects between OP- and DA-fed worms with the same genotype. * indicates comparison between different strains feeding the same diet. * ^/#^ *P* < 0.05, ** ^/##^ *P* < 0.01, *** ^/###^ *P* < 0.001, **** ^/####^ *P* < 0.0001.

To dissect how *fat-7* responds to the DA diet to regulate lipid content, we employed a forward genetic screen to identify genes that mediate the downregulation of FAT-7::GFP in the context of the DA diet (Figure 2D). We used ethyl methanesulfonate (EMS) to mutagenize the *fat-7* reporter strain, DMS303, screened ∼19,022 haploid genomes, and isolated F2 mutants with high GFP reporter intensity on the DA diet. By whole genome sequencing^35^ and EMS density mapping, we identified three mutants, *pmp-5(tp217)*, *sams-1(tp219)* and *cept-1(tp218),* that suppressed diet-mediated *fat-7* repression (Figure 2E), and verified their effect on *fat-7* transcript levels using the available mutants by qRT-PCR (Supplementary Figure S2A).

Our screen identified the 1C cycle gene *sams-1*, which converts the methionine into S-adenosylmethionine (SAM), highlighting the importance of the 1C cycle in response to DA for *fat-7* repression. Using LC-MS, we found that DA-fed worms contain more 1C cycle metabolites- including methionine, SAM and S-Adenosyl-L-homocysteine (SAH) - than OP-fed animals (Figure 2F), in agreement with previous reports^36,37^. Intriguingly, DA bacteria contain less methionine than OP (Figure 2F), suggesting that direct dietary methionine supplementation is not responsible for the observed shift in 1C metabolism. Another hit from our screen, *pmp-5* (homolog of human ABCD4), encodes a vitamin B12 transporter that transports B12 out of lysosomes^38^. In *C. elegans*, B12 is the critical cofactor for the enzymes involving in canonical propionyl-CoA breakdown and the 1C cycle^21^. Notably, DA has been reported as a B12-rich bacterium compared to OP^16^, hinting at the possibility that the DA diet might boost dietary B12 levels. In addition to *pmp-5* and *sams-1*, our screen also hit the PC synthesis gene, *cept-1*, which encodes choline/ethanolamine phosphotransferase, converting CDP-choline into phosphatidyl-choline (PC) (Figure 2E). SAM serves as the methyl group donor for converting phosphoethanolamine to phosphocholine in the PC synthesis pathway^39^, promoting us to hypothesize that PC synthesis pathway might act downstream of the B12-stimulated 1C cycle to influence *fat-7* expression.

To test the involvement of our hypothesized B12-SAM-PC pathway, we supplemented B12 to OP-fed worms. Strikingly, B12 supplementation was sufficient to suppress FAT-7::GFP expression (Figure 2G). Next, we tested each phase of the B12-SAM-PC axis by supplementing B12 to the mutants we isolated from the forward genetic screen, showing that B12-mediated downregulation of FAT-7::GFP is completely abrogated in *pmp-5*, *sams-1,* or *cept-1* mutants (Figure 2H, Supplementary Figure S2B). Moving beyond the screen hits, we employed mutants of a key enzyme of the PC synthesis pathway, *pcyt-1*^40^, and observed that these mutants were unable to repress *fat-7* mRNA expression in response to the DA diet or B12 supplementation (Supplementary Figure S2C), and had reduced PC levels under both diet regimens (Supplementary Figure S2D). Using O.R.O staining, we found that B12 supplementation further reduced lipid content in DA-fed wild-type worms (Figure 2I). This effect was suppressed in *sams-1* and *pcyt-1* mutants fed with DA (Supplementary Figure S2E, F), indicating a clear role for B12, the SAM pathway and PC synthesis in the dietary regulation of lipid (Figure 2J).

### Phosphatidylcholine synthesis as a central pathway in lipid metabolic control

Consistent with the activation of the PC synthesis pathway in DA-fed worms, we found that the total PC level - and, in particular, the level of PC32, PC34, PC36 and PC38 - is also higher in DA-fed worms compared to OP-fed animals (Figure 3A, Supplementary Figure S2D, G). To determine whether PC levels have a causal effect on organismal lipid content, we supplemented worm diets with choline, the PC precursor, and uncovered a dose-dependent decrease in lipid content as measured by O.R.O staining (Figure 3B). These data suggest that the high PC level induced by DA diet is responsible for the reduced lipid content in the worms.

**Figure 3.**
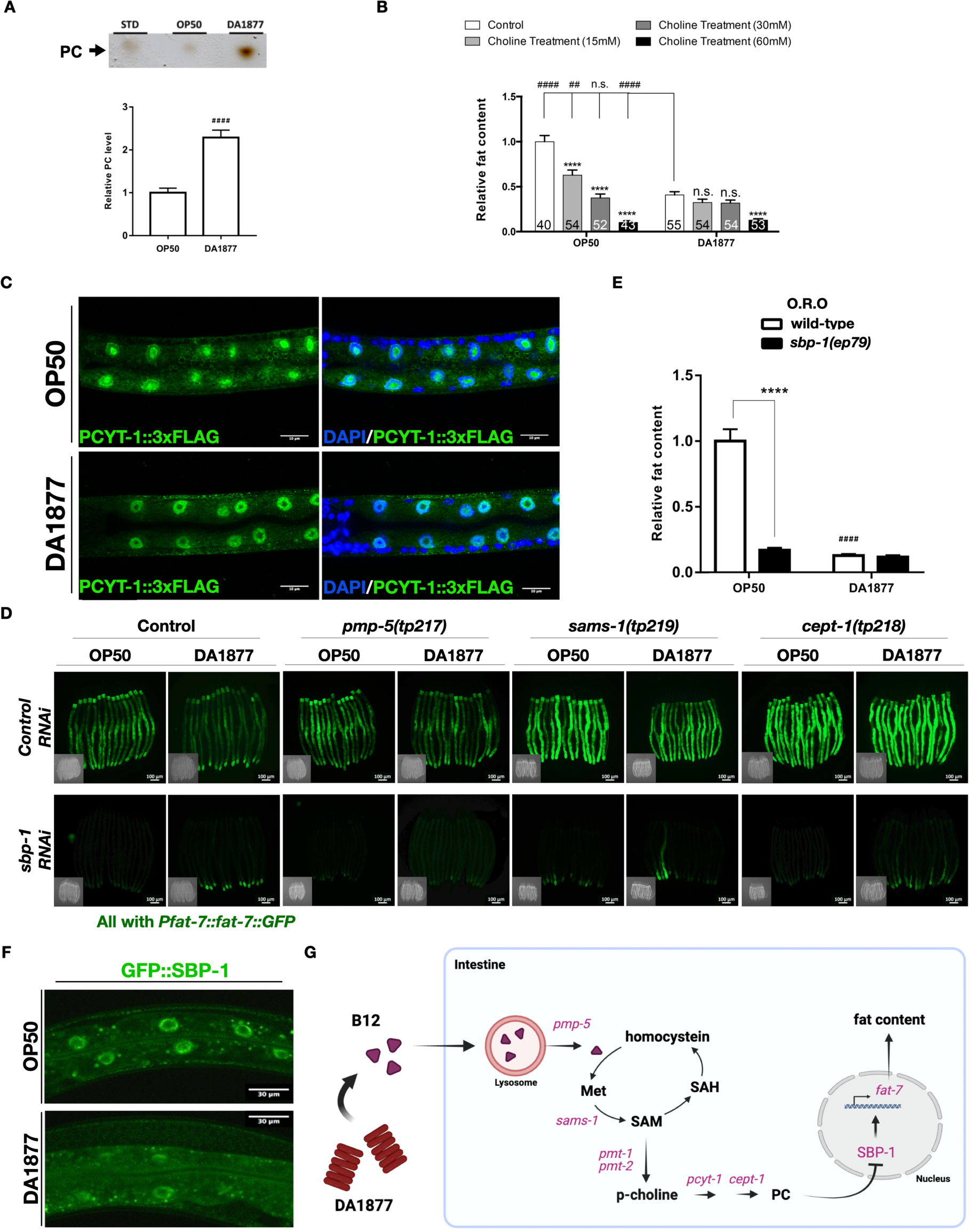
DA promotes PC level to repress lipid content in a SBP-1-dependent manner. (A) TLC analysis showing PC level in wild-type animals feeding OP or DA. (B) O.R.O staining results of wild-type worms supplemented with different concentrations of choline feeding OP or DA. Experiment contains 3 biological repeats and the number at the bottom of each graphic bar indicates the n value of each sample. (C) Representative confocal images of PCYT-1::3xFLAG (Green) in the intestinal nucleus of worms fed different diets with DAPI staining (Blue). (D) Images of FAT-7::GFP signals in the mutants isolated from forward genetic screen under *control (RNAi)* or *sbp-1 (RNAi)* treatment. (E) O.R.O staining results of wild-type and *sbp-1(ep79)* worms feeding OP or DA diet. Experiment contains 3 biological repeats and n ≥ 50 for each repeat. (F) Representative confocal image of GFP::SBP-1 localization in the intestine of worms feeding either OP or DA. (G) Cartoon for the mechanism of B12 in regulation of fat content under DA diet. This graphic is created with BioRender.com. Statistical analysis was performed using Two-tailed T tests and the data is presented as mean ± SEM. # indicates comparison of dietary effects between OP- and DA-fed worms with the same genotype. * indicates comparison between different strains feeding the same diet. * ^/#^ *P* < 0.05, ** ^/##^ *P* < 0.01, *** ^/###^ *P* < 0.001, **** ^/####^ *P* < 0.0001.

Previous studies from other species have indicated the rate-limiting PC synthesis enzyme PCYT-1 displays dynamic subcellular localization, and its catalytic activity is known to be regulated by membrane association^41^. To address whether *C. elegans* PCYT-1 localization, and therefore catalytic activity, is regulated by diet, we tagged the endogenous *pcyt-1* gene by CRISPR-Cas9 technique^42^ to observe its subcellular localization using immunofluorescence. We found that endogenous PCYT-1 localizes to the nuclear envelope and nucleolus in the intestine of OP-fed worms, but relocalizes into the nucleoplasm in response to the DA diet (Figure 3C). Taken together, these results demonstrated that the localization of PCYT-1 is differentially regulated by diets. Although the exact functional consequences of various PCYT-1 localization patterns are still unknown, it seems likely that the differential localization patterns of PCYT-1 reflect different PC demands under different dietary states. Importantly, the intranuclear localization of PCYT-1 in response to the DA diet agrees with the high PC levels that our model would suggest.

### SBP-1 is the key transcription factor downstream of B12-SAM-PC axis

We reasoned that there must be transcription factors that function downstream of the SAM-PC axis to downregulate the expression of *fat-7* and thereby lower organismal lipid content in response to DA. In *C. elegans*, *fat-7* expression has been shown to be regulated by SBP-1, NHR-49, NHR-80 and SKN-1^43–46^. In particular, low PC has been shown to promote nuclear translocation of SBP-1^47^, which prompted us to hypothesize that high PC in DA-fed worms may reduce *fat-7* expression by inhibiting *sbp-1*. We found that the FAT-7::GFP reporter expression was abolished in *sbp-1* mutants, irrespective of diet (Figure 3D), pointing to the critical role of this transcription factor in regulating *fat-7*. The increased *fat-7* expression we observed in *pmp-5, sams-1,* and *cept-1* mutants isolated from our screen was suppressed by *sbp-1* RNAi treatment (Figure 3D). Similarly, the increased lipid content in DA-fed *pcyt-1* mutants is abolished in *sbp-1, pcyt-1* double mutants (Supplementary Figure S2H). In addition, while we observed a drastic reduction of fat content in OP-fed worms lacking *sbp-1*, the DA-fed *sbp-1* mutants did not exhibit any further decrease in lipid contents (Figure 3E), indicating that DA-fed worms lack *sbp-1*-dependent lipogenesis. Intriguingly, our RNA-seq results showed that *sbp-1* was more highly expressed in DA-fed worms compared to OP-fed worms (DA/OP=1.60, *Padj* = 0.0074), implying that the reduction of SBP-1 activity is likely regulated post-translationally. Indeed, we observed less nuclear localization of a GFP::SBP-1 fusion reporter in DA-fed worms compared to OP-fed animals (Figure 3F). Together, these data suggest that the changes in lipid content that we observed across diets are orchestrated by SBP-1 and B12-SAM-PC axis mutants acts through regulation of its nuclear localization (Figure 3G).

### Diet regulates lipid droplets dynamics through SEIP-1

In addition to the changes we observed in PCYT-1 nuclear localization, we also detected PCYT-1 on lipid droplets (LDs) in OP-fed animals but not DA-fed worms (Supplementary Figure S3A). LDs store cellular lipids and are primarily present in the intestine of *C. elegans*^48^. To understand how LDs might contribute to the dietary regulation of lipid content, we examined the abundance and size distribution of LDs using confocal microscopy of the LD marker DHS-3^49^ in worms expressing a DHS-3::GFP reporter. As expected, DHS-3::GFP displayed the characteristic LD morphology in intestinal cells (Figure 4A). It is noted that the DHS-3::GFP rings were distributed more evenly in OP-fed animals, whereas clusters of rings were often observed in DA-fed animals (Figure 4A, indicated as red arrowheads). Quantitative analysis of the number and the size distribution of LDs in different diet-fed animals by image processing and analysis with MATLAB revealed that DA-fed worms possess fewer and smaller LDs compared to those fed the OP diet (Figure 4B, C). Taken together, we observe that the DA diet not only reduces total lipid content but also impacts the size and number of LDs in the intestine of *C. elegans*.

**Figure 4.**
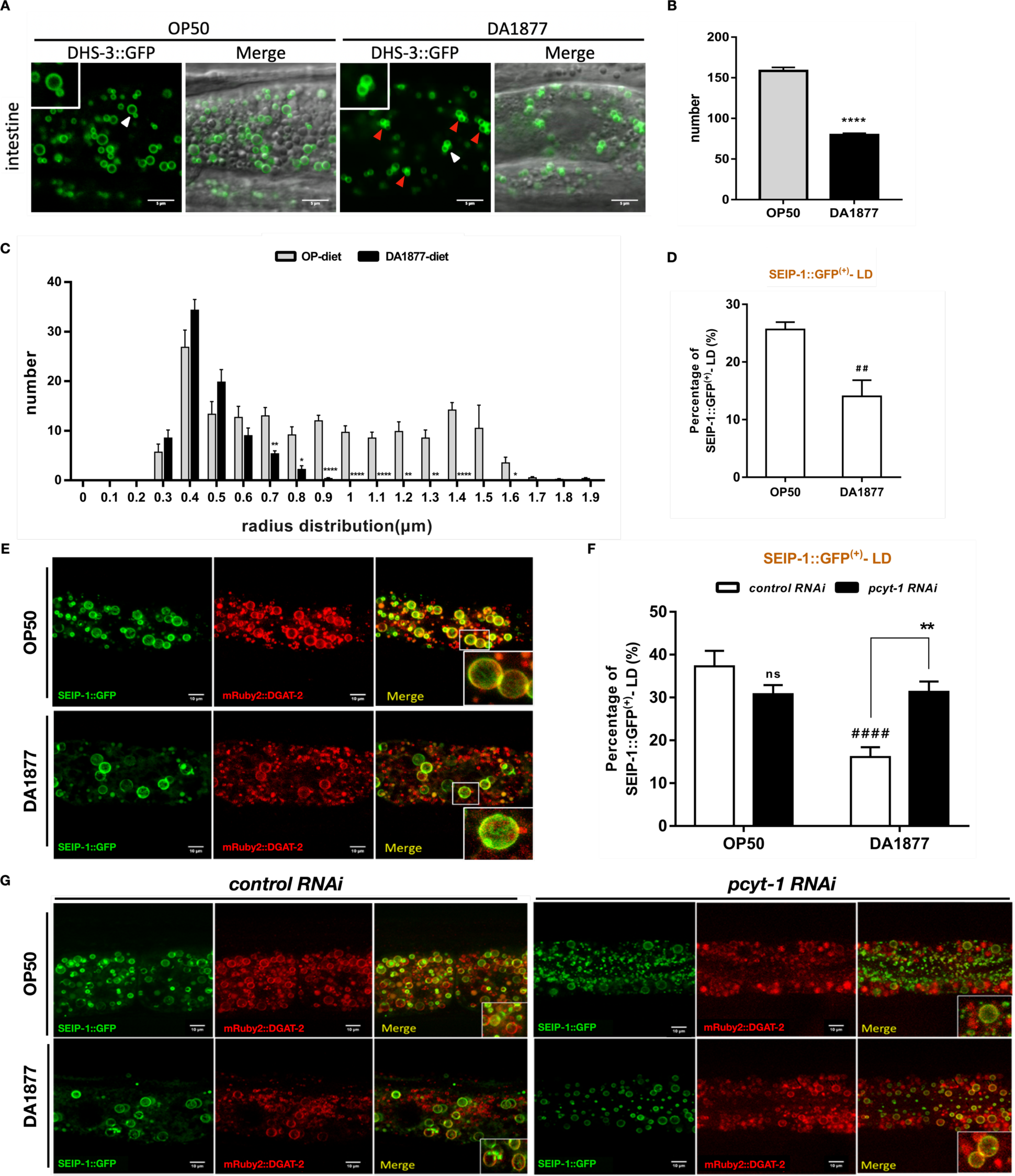
DA1877-fed worms display significantly less and smaller LDs, and have decreased SEIP-1 targeting to peri-LD cages. (A) Representative images presenting the number and size of intestinal LDs visualized by LD marker protein DHS-3::GFP in DA or OP-fed worms. Scale bar = 5μm. Insets are magnified views from white arrowheads and red arrowheads indicates the clusters of LD rings. (B, C) Quantitative results of numbers and the size of DHS-3-labeled LDs using MATLAB software. * indicates comparison between OP- and DA-fed wild-type worms. (D) Quantitative results showing percentage of LDs associated with SEIP-1::GFP in OP- or DA-fed worms. n ≥ 7 for each sample. (E) Representative images for visualization of SEIP-1::GFP (Green) in the intestine of wild-type worms. mRuby2::DGAT-2 was used as LDs marker (Red). (F) Quantitative results showing percentage of LDs associated with SEIP-1::GFP in OP- or DA-fed wild-type worms under *control RNAi* or *pcyt-1 RNAi* treatment. (G) Representative images of the intestine for visualization of SEIP-1::GFP (Green) in the *control RNAi* or *pcyt-1RNAi* treated wild-type worms feeding OP or DA. mRuby2::mDGAT-2 was used as LDs marker (Red). Statistical analysis was performed using Two-tailed T tests and the data is presented as mean ± SEM. Unless indicated, # indicates comparison of dietary effects between OP- and DA-fed worms with the same genotype. * indicates comparison between different strains feeding the same diet. * ^/#^ *P* < 0.05, ** ^/##^ *P* < 0.01, *** ^/###^ *P* < 0.001, **** ^/####^ *P* < 0.0001.

SEIP-1, the evolutionarily conserved seipin protein, controls the morphology and biogenesis of LDs, and is known to be regulated by dietary lipids^33^. This prompted us to ask whether SEIP-1 participates in the diet-mediated effects on LD dynamics we observed. Our RNA-seq analysis revealed that the DA diet decreases host *seip-1* expression (DA/OP=0.78, *Padj* = 0.015), and *seip-1* mutants have decreased total lipid content and number and size of LDs, compared to wild-type worms, on both diets (Supplementary Figure S3B, S3C). To further address how the DA diet affects the biogenesis of LDs via SEIP-1, we used mRuby2::DGAT-2 to mark the LDs^50^ and found that approximately 26% of LDs in the intestinal cells of OP-fed worms are surrounded by SEIP-1::GFP as peri-LD cages while only 14% of LDs in DA-fed young adult worms showed SEIP-1::GFP colocalization (Figure 4D, E). The reduced colocalization of SEIP-1 to LDs in response to DA diet suggests attenuation of LD emergence or subsequent expansion from the ER subdomains^33^, which agrees with our characterization of LD size and number (Figure 4A, B, C). Interestingly, the peri-LD localization of SEIP-1::GFP in DA-fed worms was boosted by RNAi-mediated knockdown of *pcyt-1,* the PC synthesis enzyme (Figure 4F, G). Together, these data reveal that the DA diet represses SEIP-1 expression, reduces its targeting to peri-LD cages in a manner dependent on the PC synthesis pathway, and thereby inhibits the biogenesis of LDs and results in decreased fat content in worms.

### Acid sphingomyelinase regulates fat content in DA-fed worms through PC

Our RNA-seq data indicated that the expression of acid sphingomyelinase (*asm-3*) – which produces Phospho-choline, a key metabolic intermediate in the PC synthesis pathways^51^ – is increased in DA-fed worms by ten-fold compared to OP-fed worms (Figure 1C). We hypothesized that ASM-3 may participate in DA-mediated fat reduction by providing phospho-choline. We first confirmed the increase in *asm-3* expression with transcriptional and translational reporters (Supplementary Figure S4A). We found that the two available mutants of *asm-3* suppressed DA-mediated fat reduction, which could be rescued by overexpression of *asm-3* (Supplementary Figure S4B, Figure 5A). Next, we analyzed *asm-3; pcyt-1* double mutants, finding that lipid content is not further increased by *asm-3* mutation in *pcyt-1* mutants upon DA feeding (Figure 5B). This suggested to us that *asm-3* acts in the same genetic pathway as *pcyt-1*. Indeed, choline supplementation was sufficient to reduce fat content in DA-fed *asm-3* mutants (Supplementary Figure S4C), supporting the upstream regulatory role of *asm-3* on the PC synthesis pathway. Interestingly, we also found higher expression of *fat-1*, *fat-2*, *fat-3*, *fat-5, fat-6,* and *fat-7* in *asm-*3 mutants, indicating that fatty acid metabolism is broadly altered in DA-fed *asm-3* mutants (Supplementary Figure S4D). Altogether, the high PC level in DA-fed worms, which impacts LD dynamics and total lipid composition, results from a convergence of SAM and ASM-3 levels.

**Figure 5.**
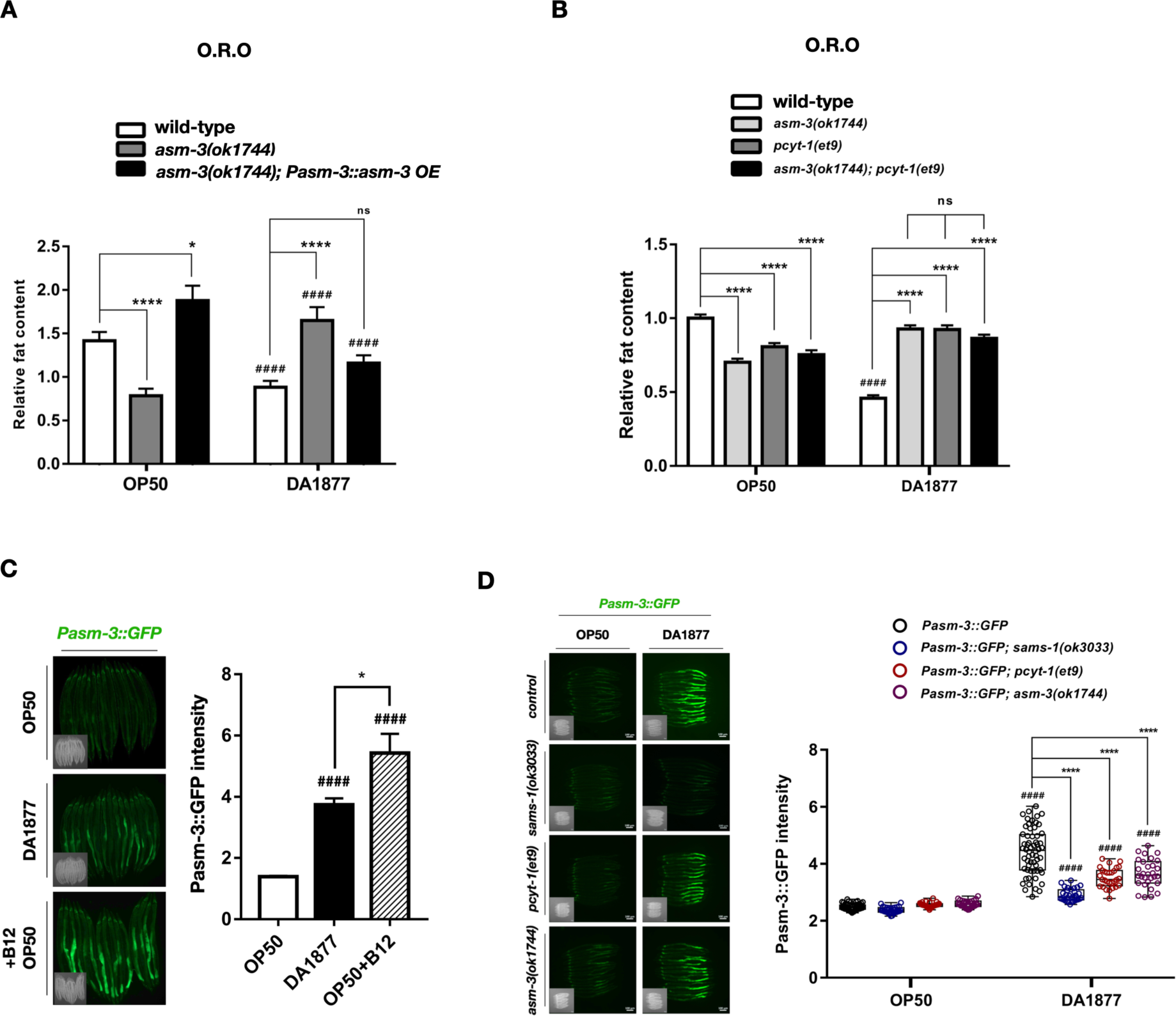
*asm-3* genetically functions together with PC synthesis pathway to regulate fat content in DA-fed worms and is transcriptionally upregulated through B12-SAM-PC axis. (A) O.R.O staining results of wild-type, *asm-3(ok1744)* mutants and *asm-3(ok1744)*; *tpEx941* mutants, which contains overexpressed *asm-3* driven by *asm-3* promoter. (B) O.R.O staining results of wild-type, *pcyt-1(et9), asm-3(ok1744),* and *asm-3(ok1744); pcyt-1(et9)* mutants. (C) Images and quantitative results of *Pasm-3::*GFP signals in worms under OP, DA, or OP with B12 supplementation diet. (D) Images and quantitative results of the signals of *asm-3* transcriptional reporter in wild-type, *sams-1(ok3033)*, *pcyt-1(et9),* and *asm-3(ok1744)* mutant background. Statistical analysis was performed using Two-tailed T tests and the data is presented as mean ± SEM. # indicates comparison of dietary effects between OP- and DA-fed worms with the same genotype. * indicates comparison between different strains feeding the same diet. * ^/#^ *P* < 0.05, ** / ## *P* < 0.01, *** ^/###^ *P* < 0.001, **** ^/####^ *P* < 0.0001.

This prompted us to ask whether *asm-3* expression is regulated by the B12-SAM-PC axis. We found that supplementation of B12 to OP-fed worms was sufficient to drive expression of the ASM-3::GFP reporter or endogenous *asm-3* to higher levels as observed in DA-fed worms (Figure 5C, Supplementary Figure S5A). *asm-3* expression was significantly reduced in B12-SAM-PC axis mutants (*pmp-5(tm5497)*, *sams-1(ok3033), cept-1(et10),* and *pcyt-1(et9)* (Figure 5D, Supplementary Figure S5B, C). Since ASM-3 participates in PC synthesis pathway by producing the intermediate metabolite, phospho-choline, these results suggest *asm-3* may auto-regulate itself. Indeed, we found that *asm-3* expression is decreased in *asm-3* mutants under DA feeding when compared to wild-type worms (Figure 5D). Furthermore, we found that *asm-3* expression is slightly increased in *sbp-1(ep79)* mutants feeding DA (Supplementary Figure S5D).

Altogether, these results reveal the importance of the B12-SAM-PC axis for regulating *asm-3* expression, and suggest that *asm-3* constructs a positive feedback loop to potentiate PC levels. Using an ASM-3::mCherry reporter, we noted that ASM-3 protein is detectable not only in intestinal cells, exclusively in the middle and posterior sections of the intestine (Supplementary Figure S4A), but also in scavenger coelomocytes (Figure 6A). Specifically, by introducing a lysosomal marker, LMP-1::GFP^52^, we found that ASM-3 is localized in the lysosomes of coelomocytes (Figure 6B). Human ASM is known to be expressed in both an endolysosomal (L-ASM) and secretory (S-ASM) form^53^, and we wondered whether secretion of intestinally-expressed ASM-3 could be the source of the observed coelomocyte ASM-3 localization. Using the intestine-specific *vha-6* promoter to drive *asm-3* expression, we were still able to detect ASM-3 in intestinal cells, pseudocoelomic fluid, and coelomocytes (Figure 6C, Supplementary Figure S6A), indicating that *asm-3* is transcribed and translated in the intestine, and later enters pseudocoelomic fluid and coelomocytes by secretion. To decipher whether ASM-3 functions in coelomocytes to regulate fat content, we expressed *asm-3* using the coelomocyte-specific *unc-122* promoter^54^ (Supplementary Figure S6B), or the intestine-specific *vha-6* promoter. Strikingly, coelomocytes-restricted or intestine-restricted expression of *asm-3* were both sufficient to reduce fat content in *asm-3* mutants (Figure 6D). These results suggest that ASM-3 may signal from the intestine to coelomocytes to regulate *C. elegans* lipid content in response to diet. Together, our data pointed out the importance of the coelomocyte localization of ASM-3 on fat regulation under high B12 diet in worms.

**Figure 6.**
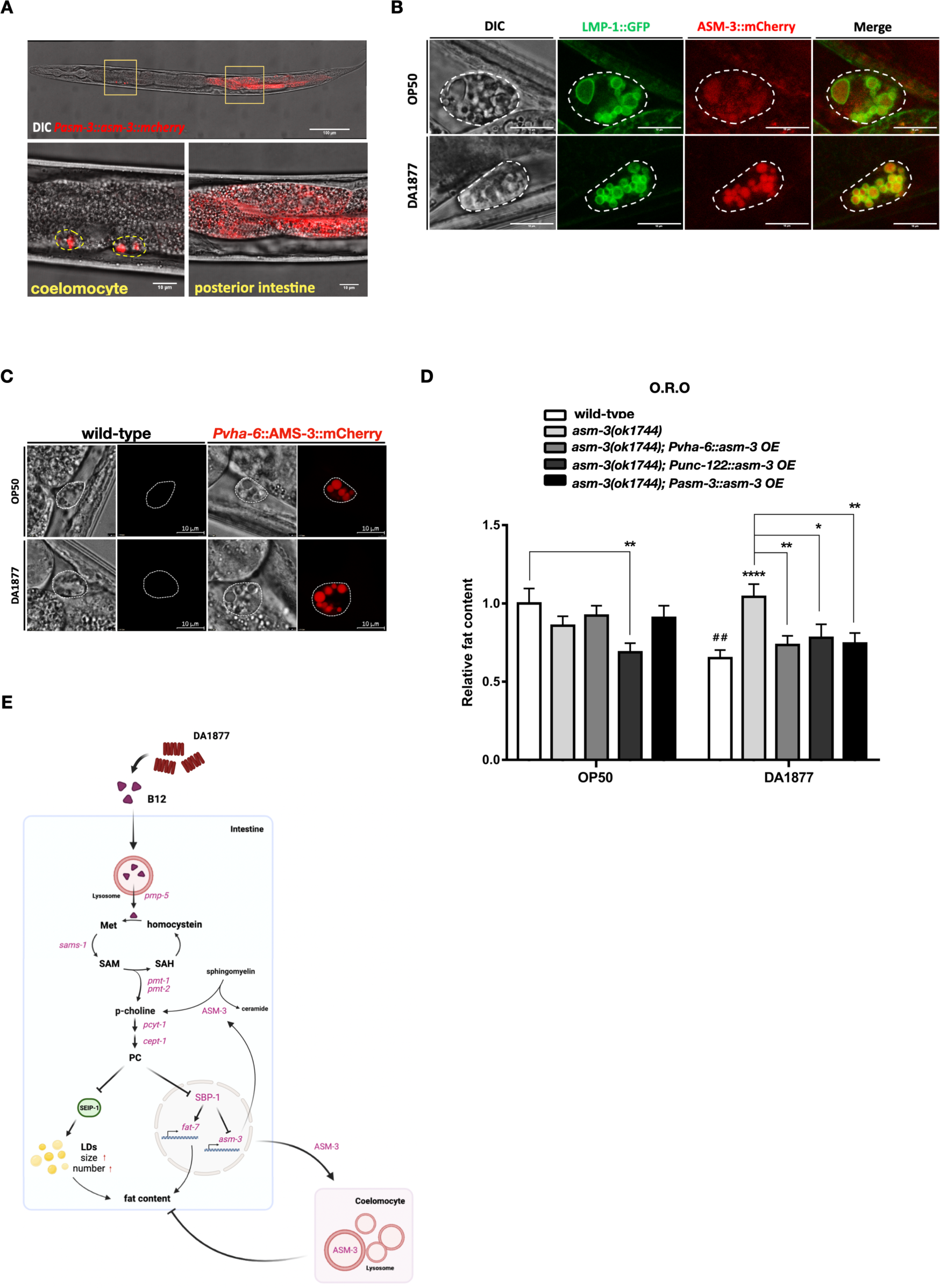
ASM-3 transports to coelomocytes from intestine to regulate fat content in DA-fed worms. (A) Representative images revealing the localization of ASM-3::mCherry (Red) in YW2989 [*tpEx941*] worms where ASM-3 was driven by its own promoter. (B) Images of ASM-3 localization in the coelomocyte of YW3735 strain, which contains single-copy insertion of *asm-3::mCherry* (Red) and lysosomal marker LMP-1::GFP (Green). White dotted lines mark coelomocytes. Scale bar = 10µm (C) Images presenting the localization of intestine-specific expressed ASM-3::mCherry (Red) in YW3034 [*asm-3(ok1744); tpEx944*]. White dotted lines mark coelomocytes. (D) O.R.O staining results of wild-type worms, *asm-3(ok1744)* mutants, *asm-3(ok1744)* with intestinal *asm-3* expression using intestine-specific promoter *vha-6, asm-3* expression in coelomocytes using coelomocyte-specific promoter *unc-122,* or *asm-3* expression driven by the endogenous *asm-3* promoter. Statistical analysis was performed using Two-tailed T tests and the data is presented as mean ± SEM. # indicates comparison of dietary effects between OP- and DA-fed worms with the same genotype. * indicates comparison between different strains feeding the same diet. * ^/#^ *P* < 0.05, ** ^/##^ *P* < 0.01, *** ^/###^ *P* < 0.001, **** ^/####^ *P* < 0.0001. (E) Model showing mechanism of how DA1877 diet regulates fat content in the worms. This graphic is created with BioRender.com.

## Discussion

Vitamin B12, although required in minute quantities, plays a crucial role as an essential micronutrient that significantly impacts human health^55^. Previous studies from *C. elegans* have illuminated that elevated dietary B12 accelerates 1C cycle activity and reroutes propionate breakdown, thus driving host metabolic plasticity^16,21^. Our study corroborates and extends this understanding by pinpointing a Δ9 fatty acid desaturase, *fat-7,* as a key factor and revealing the intricate mechanisms through which B12 regulates lipid homeostasis. These results offer valuable insights that may inform research and therapeutic strategies for metabolic disorders by targeting B12-impacted cellular pathways.

Our study showed that the DA diet or B12 supplementation promotes the levels of PC, which in turn exert multifaceted effects on fat regulation, such as modulating lipogenic gene expression via SBP-1 (Figure 3D-G) and affecting LD budding and expansion via SEIP-1 (Figure 4D-G). Interestingly, PC is known to be one of the primary components of lipoprotein^56^ and is critical for very-low-density lipoprotein (VLDL) secretion^57–59^. This suggests the intriguing possibility that high PC may also promote TAG export. In circumstantial support of this notion, our RNA-seq analysis indicated that *dsc-4* (the *C. elegans* ortholog of the large subunit of the microsomal triglyceride transfer protein (MTP)^60^ and five apoB-like genes ( *vit-2*, *vit-3, vit-4, vit-5,* and *vit-6*^61^ ) are upregulated in DA-fed worms compared to OP-fed ones (data not shown), suggesting the possibility that the secretion of TAG may be highly activated in response to the DA diet. Further work will be required to elucidate the effects of dietary B12 on TAG export through the lipoprotein pathway, and whether this depends on PC.

In addition to the regulatory effect that PC exerts on LD biogenesis, previous work has also implicated cyclopropane fatty acids (FAs) and specific PUFAs in SEIP-1 enrichment at peri-LD cages^33^. Our LC-MS results indicated that cyclopropane FA levels are undetectable in DA bacteria (Figure 1G). Since *C. elegans* are unable to synthesize cyclopropane FAs and rely on dietary sources for these compounds^62,63^, DA-fed worms have a lower abundance of cyclopropane FAs, in addition to reduced levels of most PUFAs, compared to OP-fed worms (Figure 1F). We therefore speculate that the decreased targeting of SEIP-1 to peri-LD cages in DA-fed worms may be partly explained by the lower levels of cyclopropane FAs and PUFAs. Further, *C. elegans* use cyclopropane FAs for a substantial portion of the TAGs they produce^63^. It is therefore possible that lower levels of cyclopropane FAs in DA-fed worms may also impede TAG synthesis, resulting in overall reduced lipid content (Figure 1A, B). Together, these results suggest the possibility that the altered lipid homeostasis we observed, while principally driven by the high B12 level provided by DA, may also be influenced by other dietary components like cyclopropane FAs and PUFAs contributed by DA.

*sams-1*, responsible for the conversion of methionine to SAM, is upregulated in worms fed DA, leading to higher SAM levels (Figure 2F). SAM serves as a universal methyl donor, providing a methyl group for PC synthesis, DNA, and histone methylation^39^. Loss of *sams-1* significantly increases fat content by more than three-folds (Supplementary Figure S2E); given its pleiotropic role in many metabolic pathways, this suggests that *sams-1* likely modulates fat content in DA-fed worms through multiple pathways beyond the PC synthesis. Indeed, this is further supported by our observation that mutants of histone methyltransferases SET-2 and SET-30 (which install H3K4me1 and H3K4me2, respectively) exhibited elevated lipid content on DA diet (data not shown). Notably, while both *set-2* and *set-30* mutants had this effect, only *set-2* appears to control *fat-7* expression (data not shown), implying a vast network of lipid metabolism regulatory mechanisms governed by SAM.

Through our implication of *asm-3* coelomocyte activity in regulating lipid content, our findings established a role for coelomocytes in diet-mediated lipid homeostasis. Earlier studies have indicated the engagement of coelomocytes in starvation-induced fat mobilization and lifespan extension^64^. Together, these findings support the notion that coelomocytes can integrate nutritional cues to influence fat metabolism. We also provided evidence for inter-tissue communication of ASM-3 signaling from expression in intestinal cells to action in coelomocytes, opening up new avenues of research into intercellular communication in the dietary control of metabolism.

Altogether, our results reveal that dietary vitamin B12, enriched in the DA bacteria diet, stimulates the SAM-PC axis to inhibit lipogenesis in a *sbp-1*-dependent manner and upregulate *asm-3* expression, together resulting in reduced lipid content and reduced LD biogenesis in *C. elegans* (Figure 6E). The diet-induced metabolic rewiring underscores the complex networks of cellular and organismal metabolic control, highlights the major effect of micronutrients on primary metabolism, and paves the way for future insights into dietary supplementation and metabolic disease.

## Materials and Methods

### *C. elegans* culture and strains

Worms were grown and maintained at 20°C on standard nematode growth medium (NGM) plates seeded with bacteria. The Bristol N2 strain was used as wild-type. A list of mutants and transgenic strains is available in the Key Resources Table.

### Bacterial strains and culture conditions

*E. coli* OP50 and *C. aquatica* DA1877 were used as the food for worms. Bacterial cultures were prepared from a single colony in 5ml LB medium (Sigma, L3022) with shaking at 37°C for 16 hours, and then sub-cultured into 50ml LB until the OD600 reaches 1.5 to 1.8. Subsequently, 200 μL of OP50 or DA1877 were spread on a 6 cm NGM plate and kept at RT for two days to let bacteria grow and dry before use.

### Plasmid construction

Molecular cloning was mostly conducted by Multisite Gateway Three-Fragment Vector Construction system (Invitrogen, #12537-023). The plasmid pYW1693 (*Pvha-6::fat-7::eGFP_let-858_3’-UTR*) was made by recombination of pML25 containing 0.9kb *vha-6* promoter, *fat-7* open reading frame (ORF) and pGH112 containing *eGFP_let-858_3’-UTR*. *fat-7* ORF was amplified using genomic DNA from the start codon to the end of *fat-7* gene before the stop codon by the following primers: attB1_fat-7_f / attB2_fat-7_r, and cloned into pDONR221 through BP reaction. The plasmid pYW1783 (*Pasm-3::gfp_let-858_3’UTR*) was constructed by combination of pCR110 containing GFP with ATG, pADA126 containing *let-858_3’UTR* and *asm-3* promoter entry clone including 2.6kb of *asm-3* promoter region amplified using primers: asm-3p_f / asm-3p_r, and cloned into pDONRP4-P1R. The plasmids pYW1784 (*Pasm-3:asm-3::mCherry_let-858_3’-UTR*), pYW1850 (*Pasm-3::asm-3::mCherry_let-858_3’UTR*), pYW1785 (*Pvha-6::asm-3::mCherry_let-858_3’-UTR)* and pYW1851 (*Punc-122::asm-3::mCherry_let-858_3’UTR*) were constructed by combination of gwYW37 and pCFJ159 together with promoter entry clone containing above-mentioned *asm-3* endogenous promoter, intestine-specific promoter pML25, or coelomocyte-specific promoter *Punc-122*^54^. For coelomocyte-specific promoter entry clone: 0.2kb of *unc-122* promoter region was amplified with primers: unc-122p_f / unc-122p_r and cloned into pDONRP4-P1R. gwYW37 contains the genomic region of *asm-3,* which was amplified from the start codon to the end of *asm-3* before stop codon with primers: asm-3_f / asm-3_r, and cloned into pDONR221. The vectors for pYW1693, pYW1783, pYW1784, and pYW1785 were assembled into pDEST R4-R3 Vector whereas the vectors for pYW1850 and pYW1851 were assembled into pCG150_phiC31_V2 which was kindly provided by John Wang Lab. pML25, pGH112, pCR110, pADA126, and pCFJ159 containing *mCherry_let-858_3’-UTR* was kindly provided by Erik Jorgensen Lab.

The plasmid pYW1808 is constructed to generate the dsDNA donor for CRISPR-Cas9-based *gfp::sbp-1* worms. The insert was amplified using genomic DNA from the transgenic strain CE548 [*sbp-1(ep79); epEx141(Psbp-1::GFP::sbp-1+rol-6(su1006)*] with the primers: sbp-1-CrisprA / sbp-1-CrisprB, and cloned into T&A^TM^ using T&A^TM^ cloning kit (YB Biotech, FYC001-20P).

### EMS mutagenesis and mutation identification

The whole genome sequencing (WGS)-based mutation identification strategy was followed according to Zuryn *et al*. 2010 ^35^. In brief, DA-fed L4 worms were collected into 15 mL centrifuge tubes and washed with ddH_2_O for three times to reduce bacterial contaminants. The worms were then incubated in 50mM Ethyl methanesulfonate (EMS) (Sigma, M0880) under room temperature (RT) with constant rotation for four hours. After EMS treatment, the worms were recovered for two hours. Then the recovered worms were placed on new large DA1877-seeded NGM plate (P0s, two worms on each plate). After 16 hours, P0s were removed from the plates, and their offspring were incubated under 20 °C until F2 progenies (F2s) reached adulthood. F2s with high GFP intensity were selected and cloned. The F3 progeny were scored to remove the false positives. Then, the isolated mutants were subjected to complementation tests. The endogenous *fat-7* mRNA level of the mutants was further validated using qRT-PCR. Next, the selected mutants were backcrossed to the non-mutagenized parental strain, DMS303, for 4-5 times to remove the background mutations.

For mutation identification, recombinants were collected after the backcross processes and their genomic DNA (gDNA) was extracted by using DNeasy® Blood and Tissue kit (Qiagen, #69504) following the manufacturer’s protocol. The gDNA libraries were constructed by using Illumina TruSeq® Nano DNA Library Preparation kit (#FC-121-9010DOC) according to the manufacturer’s protocol, with two amplification cycles. The paired-end WGS (2 x 150 bp) was performed on a NovaSeq 6000 system (Illumina). The bioinformatics analysis of the WGS-based mutation identification was performed according to the MiModD pipeline v0.1.9^65^ by Center for Computational and Systems Biology and Technology Commons at College of Life Science, National Taiwan University. The raw FASTQ sequencing read files were first processed with Trimmomatic v0.39^66^ for adaptor trimming and quality control. WGS reads were aligned to the WBcel235/ce11 reference genome using SNAP v0.15.4 ^67^. Next, SNP and short sequence indel calling was performed by running the MiModD script, which used the samtools and bcftools^68^ internally, on the aligned reads. Then, the called variants were annotated by using SnpEff v4.3t^69^. The variance annotation was performed based on the Ensembl release 102 gene annotation file.

### Oil Red O staining and quantification

Oil Red O (O.R.O) staining was conducted as previously described with some modification^70^. In brief, O.R.O solution (Sigma, O1391) was pre-filtered by 0.22 um filter and rocked on platform shaker, then diluted to 60% with water and placed on shaker overnight at room temperature. Next day, young adult worms were harvested into 1.5 mL tubes using S buffer (0.1 M NaCl, 0.0065 M K_2_HPO_4_, 0.0435 M KH_2_PO_4_) and washed 3 times to reduce bacterial contamination. 500 μL of 60% isopropanol was added to fix the worms and let them settle to the bottom by gravity in order to remove the supernatant. Subsequently, 500 μL of 60% O.R.O solution, which was filtered again using 0.22 um filter to prevent precipitation before adding into worm samples, was added into worm pellet. After gentle inversion, worm samples were placed in a wet chamber away from light on a shaker at 25°C for 8-10 hours. Then, the O.R.O solution was substituted by 250 μL of S buffer with 0.01% Triton-X 100 and the samples were stored at 4°C. Worms were mounted on glass slides directly and the images were captured by Zeiss Axioplan microscopy within three days. For images analysis, Image J^71^ was used to separate color channels and O.R.O intensity is measured by subtracting green channel signal from red channel signal. Statistical analysis was performed using the Student T test or One-Way ANOVA with Tukey correlation.

### RNA-seq analysis

Total RNA was extracted from the young adult worms fed either DA or OP, each with biological replicates (n = 3), using TRIzol® (Invitrogen, #15596018) following the manufacturer’s protocol. RNA integrity was assessed using Agilent 2100 Bioanalyzer (RNA 6000 Pico assay), and the concentration was measured using the Qubit device. The RNA-seq libraries were prepared using the Illumina TruSeq® Stranded mRNA kit according to the manufacturer’s protocol starting with 1 μg total RNA. The paired-end RNA sequencing (2 x 150 bp) was performed on a HiSeq 4000 system (Illumina). The sequencing data were converted from base call files to FASTQ files using Illumina’s bcl2fastq v2.2.0. The raw FASTQ files were first processed with Trimmomatic v0.39^66^ for adaptor trimming and quality control. The *C*. *elegans* genomic reference sequence WBcel235/ce11 was downloaded from Wormbase^72^, and the gene annotation file (release 100) was downloaded from Ensembl^73^. The paired-end reads were aligned and mapped to *C*. *elegans* genomic reference sequence using HISAT2 v2.2.0^74^, followed by quantifying the raw read counts of individual genes using featureCounts v2.0.1^75^. The differential expression between different diet-fed worms was statistically assessed using the R/Bioconductor package DESeq2 v1.28.1^76^. Genes with a Benjamini-Hochberg adjusted *p*-value < 0.01 were defined as differentially expressed genes (DEGs). Gene Ontology (GO) and KEGG pathway enrichment analyses of DEGs were performed by DAVID^77^.

### Lipidomic analysis

Synchronized young adults were washed off of NGM plates into 15 ml centrifuge tube using M9 buffer. The worm pellet was then washed with M9 buffer for 3 times of to remove bacteria. After aspirating the supernatant, distilled water was added to the worm pellet and transferred into a 1.5 ml tube. Next, the worm pellet was washed using 30mM NaN3 for three times and collected into the homogenizer tube with 100 μL of Zirconia beads (BioSpec, #11079110zx), followed by immediate immersion in liquid nitrogen for at least 1 hour. The samples were homogenized using Precellys Evolution (Bertin) with a program setting containing three cycles of 6400 rpm 20 seconds, 10 second rest and 6400 rpm 20 seconds again; whereas for bacteria OP50 or DA1877, samples were transferred into VK01 microorganism lysis kit (Thermo Fisher Scientific) and homogenized with a program setting consisting of 10 cycles of 9500 rpm for 30 seconds, rest in machine for 60 seconds, 9500 rpm for 30 seconds and rest on ice for 60 seconds. After homogenization, samples were centrifuged at 13000 rpm for 10 mins (worms) or 4000 rpm for 20 minutes (bacteria) at 4°C. Worm lysates were transferred into glass tube and a small portion of lysate was used to measure the concentration of total protein by BCA^TM^ Protein Assay Kit (PIERCE). Next, 2 mL ice-cold (-20°C) chloroform-methanol mixture (2:1, v/v) and 20ul of the internal control (FA13:0) were added to samples for lipid extraction. First, samples were vortexed and incubated at 4°C overnight on shaker. Next day, samples were split into two 1.5 mL tubes and then 400 μL of Hajra’s solution (0.2 M H_3_PO_4_ and 1 M KCl) was added to each tube, followed by vortexing to ensure good mixing. After centrifugation at 2000rpm for 10 mins, the aqueous (upper) and organic layer were separately transferred into new tubes, respectively. 750 μL of chloroform were added to the aqueous layer, followed by centrifugation at 2000 rpm for 10 minutes to re-separate the organic layer. Next, this chloroform layer was pooled with the previous organic layer. The samples were dried by SpeedVac (Thermo Scientific), then the tube was filled with argon gas to store the samples at -20°C. Chloroform/methanol (2:1) was added to dissolve the sample before use.

For LC-MS analysis, we used Orbitrap Elite Ion Trap-Orbitrap mass spectrometer (Thermo fisher Scientific) coupled with a ACQUITY UPLC/UHPLC system (Waters). The MS was operated with either positive or negative ion modes using the full Fourier transform-mass spectrometry scan at 100-1200 m/z, resolution 60,000. The phospholipid and TAG species were identified by LC/MS/MS (personal communication with Chao-Wen Wang Lab). Lipid abundance was quantified with Xcalibur software (Thermo Scientific).

To analyze fatty acids with GC-MS, 230 μL of 14% boron chloride-methanol was added into dried lipid extraction^78^ to prepare for the fatty acid methyl esters (FAMEs). After 20 minutes incubation for complete dissolution, the samples were heated to 95°C in a water bath for 10 minutes. After the sample were cooled down to room temperature, 200 μL of benzene were added and samples were heated again to 95°C for 30 minutes. Next, samples were cooled to RT again for 10 minutes, followed by adding 230 μL of water and 690 μL of petroleum. The samples were vortexed and centrifuged for 3 minutes at 2000 rpm for phase separation. The organic layer (upper layer) was then transferred into a new tube and dried using SpeedVac (Thermo Scientific). For GC-MS, Agilent GC/MSD 5975C-PAL RTC120 equipped with Agilent DB-5MS column was used, 1 μL of FAME sample was injected at the split ratio of 1:10. Oven temperature was held at 80°C for 1 min, increased by 8°C /min to 128°C, and increased by 10°C /min to 188°C /min, increased by 2°C /min to 222°C, increased by 3°C /min to 228°C , increased by 5°C/min to 278°C. The fatty acids quantification results of each sample were normalized to its own internal control level, FA13:0, which was added into worm lysate before lipid extraction, and their respective protein level. The fatty acids quantification results of bacterial samples were normalized to their OD600 measured at collection.

### Metabolites extraction

Synchronized worms were collected into a homogenization tube with 200 μL of 50% methanol and 100 μL of 1 mm Zirconia beads (BioSpec, #11079110zx). Worms were homogenized by Precellys Evolution (Bertin) with program setting as three cycles of 6400 rpm 20 seconds, 10 second rest and 6400 rpm 20 seconds again. Of note, the samples were incubated on ice for 10 seconds between cycles. After homogenization, the samples were centrifuged at 10000 rpm for 10 minutes at 4°C and supernatant was then transferred into a new 1.5 mL tube. Each sample was deproteinized by adding 200 μL of 31% acetonitrile, which was dissolved in formic acid. The samples were vortexed and cooled on ice for 5 minutes. The samples were air-dried at room temperature and then the pellets were solubilized with 500 μL of methanol. The samples were filtered into a new tube using 0.22 μm PVDF syringe filter and stored at 4°C before analysis. For extracting metabolites from bacteria, we added 2 mL methanol into the collected pellet of OP50 or DA1877 and transferred the samples into glass tubes. Next, 1 mL chloroform was added and samples were mixed by vortex. Samples were incubated at room temperature for 30 minutes and placed on a shaker at 4 °C overnight. The next day, the sample was aliquoted into three 1.5 mL tubes before addition of 400 μL of Hajra’s solution (0.2 M H_3_PO_4_ and 1 M KCl in 10 mL ddH_2_O). The aqueous and organic layers were separated by centrifuging at 2000 rpm for 10 minutes. The aqueous layer was transferred into a new tube and dried by SpeedVac (Thermo Scientific). The pellet was solubilized with 500 μL of methanol and filtered into a new tube using 0.22 μm PVDF syringe filter. Samples were stored at 4°C before analysis.

### Thin layer chromatography (TLC) analysis

For triacylglycerol (TAG) quantification, two TLC chambers were pre-equilibrated overnight by two solvent systems: one contained solvent 1 (petroleum ether: diethyl ether: acetic acid = 70: 30: 2, v/v/v) and the other contained solvent 2 (petroleum ether: acetic acid = 49: 1, v/v). The TLC Silica gel 60G F_254_ 25 Aluminum sheets 20 × 20 cm plate (Merck, #100390) was activated by heating at 95°C for 15 minutes and then air cooled. 20 μL of lipid extraction was spotted onto the TLC plate and dried the lipid spots for 10 minutes. The plate was dipped into solvent 1 until the front of solvent 1 reached two-thirds of the height of the plate. After drying the plate for 10 minutes, it was dipped into solvent 2 in the same direction until the solvent 2 front reached 0.5 cm from the top of the plate. For visualization, the TLC plate was incubated for 10 seconds in MnCl_2_ solution (0.63 g MnCl_2_·4H_2_O, 60 ml of water, 4 mL of sulfuric acid and 60 mL of methanol), then dried for 15 minutes at room temperature and then heated at 170°C for 15 minutes. The plate image was scanned and the intensity of TAG was quantified using ImageJ. 1,2,3-oleoyl-glycerol was used as a standard (Sigma). For phosphatidylcholine quantification, two TLC chambers were pre-equilibrated overnight with solvent 3 (chloroform: ethanol: water: triethylamine = 30: 35: 7: 35, v/v/v/v) and solvent 4 (chloroform: methanol = 1: 1, v/v), respectively. The TLC plate was first dipped into solvent 4 until solvent 4 developed to the top of the TLC plate. After drying the plate for 10 minutes, it was completely wet with 1.8% boric acid solution (in 100% ethanol) for 15 seconds. The plate was then dried for 10 minutes again and then heated at 95°C for 15 minutes, before loading the samples and standard. The lipid spots were dried for 10 minutes and then the plate was dipped into solvent 3 until the solvent 3 front reached two-thirds of the height of the plate. The plate was then dried for 10 mins before continuing the separation in solvent 3 for another 1 hr 45 mins. The plate was then dried overnight. To visualize phosphatidylcholine, the plate was soaked in 250ml solution of 8% of phosphoric acid (w/v), 10% of copper sulphate (w/v) followed by 1 hour of air-drying and 1.5 hours incubation at 170°C until the signals are clear. The intensity of phosphatidylcholine was quantified through ImageJ and 1,2-dioleoyl-sn-glycerol-3-phosphocholine (AvantiPolar Lipids) was used as a standard.

### Quantitative RT-PCR

10 worms were washed in a drop of H_2_O and collected into a PCR tube containing 2 μL of lysis buffer (50 mM KCl, 10 mM Tris pH 8.3, 2.5 mM MgCl_2_, 0.45% NP-40, 0.45% Tween-20, 0.01% Gelatin and 100 μg/mL protease K). Worms were then lysed by incubation at 65°C for 10 minutes and followed by 1 minute of 85°C to activate and inactivate proteinase K. Next, cDNA was synthesized following manufacturer’s protocol of RevertAid H minus First Strand cDNA Synthesis Kit with random hexamer primers (Thermo Scientific, #K1632). Quantitative real-time PCR was performed with iQ SYBR® Green Supermix (Bio-Rad, #1708886) using the Bio-Rad CFX96 System or CFX384. Primers used in qRT-PCR were designed to span exon-exon junctions and their sequence are listed in the primer Table. *act-4* was used as the internal control. The primers’ melting curves were examined carefully to ensure primer specificity. Relative expression levels of transcripts were calculated with the comparative CT method as previously described^79^.

### Microscopy and Quantification of Fluorescence

Worms were mounted on a 4% agar pad and anesthetized with 30 mM NaN_3_. For FAT-7::GFP, AXIO Imager.M2 (ZEISS) was used to acquire images and the GFP signals were analyzed by ImageJ. For DHS-3::GFP, SEIP-1::GFP, DGAT-2::mRuby, PCYT-1::3xFLAG and ASM-3::mCherry, the images was captured by Zeiss LSM780 or Leica SP8 confocal microscopic systems. For LDs analyses, Z-stack sectioning for at least 5 images at 2 um intervals were acquired and analyzed using MATLAB (MathWorks). Briefly, the images were first adjusted to enhance the intensity and eliminate noise. Next, the images were converted to binary with proper threshold and circle objects on images were detected and analyzed. For GFP::SBP-1, Leica Stellaris 8 confocal microscopy was used and Tau-gating function is applied to reduce autofluorescence interference in the intestine.

### Immunofluorescence staining

Immunofluorescence staining was performed using the whole-mount fixation method previously described ^80,81^ with formaldehyde solution (Sigma, F8775). Primary antibodies used were anti-FLAG (Sigma, F3165) at 1:250 and anti-GFP (abcam, ab6556) at 1:200. Secondary antibodies were anti-mouse Alexa Fluor 488-conjugated secondary antibody (Invitrogen A11001), anti-mouse Rhodamine Red-X (RRX)-conjugated secondary antibody (Jackson ImmunoResearch, 715-296-150), anti-rabbit Fluorescein (FITC)-conjugated secondary antibody (Jackson ImmunoResearch, 711-095-152), and all were used at 1:200. Samples were mounted using VECTASHIELD Plus antifade mounting media (Vector Labs, H-1900) with DAPI at the final concentration of 2ug/ml. The images were acquired using Zeiss LSM780 Confocal Microscopy System and processed using Image J^71^. Identical exposure time and setting for brightness and contrast adjustment were applied for the same set of experiments.

### RNA interference

RNA interference (RNAi) was performed by feeding worms with bacteria expressing dsRNA as described previously^82^ with some modification. In brief, the NGM plates for an RNAi experiment contains 25ug/ml ampicillin and 1 mM IPTG. RNAi clones were obtained from the Ahringer RNAi library^83^ or by cloning the genomic region of the target gene into RNAi vector L4440, followed by transformation of the RNAi-compatible OP50(xu363) strain^84^ and culturing in LB medium with 25ug/ml ampicillin until the OD600 reached 1.5 to 1.8. Then, 200 μL of the bacteria was spread onto each RNAi- NGM plate and left at RT for two days before use. For DA-fed RNAi knockdown experiments, we cultured DA1877 to the OD600 of 1.5 to 1.8 and then homogenized DA1877 by Precellys Evolution (Bertin) using the VK01 microorganism kit (Thermo Fisher Scientific) with 10 cycles of 9500 rpm for 30 seconds, resting for 60 seconds, 9500 rpm for 30 seconds and resting on ice for 60 seconds. The DA1877 lysate was mixed with RNAi clone-containing OP50*(xu363)* at a ratio of 1:1. 400 μL of the mixture were seeded on each RNAi- NGM plate.

### B12 and Choline supplementation

Choline Chloride (Sigma, 26978) was freshly dissolved in ddH2O at a concentration of 3 M and sterilized through a 0.22μm filter. The choline solution was then added into autoclaved NGM at a final concentration of 15 mM, 30 mM or 60 mM. Choline plates were placed at RT overnight and seeded with bacterial culture before use. The worms were cultured on choline-supplemented NGM plates for 3 generations before analysis.

Vitamin B12 was dissolved in ddH2O to make a stock solution of 64mM (Sigma, V2876). The B12 stock was then diluted to 32nM with water and 200ul were spread on each 6 cm NGM plate with seeded bacteria. The plates were allowed to dry for 30 mins in a laminar flow hood until the B12 solution was completely absorbed by NGM. Notably, B12 is photodegradable and therefore all B12 experiments were performed in the dark.

### CRISPR-Cas-9 Knock-in

The CRISPR-Cas-9 genomic knock-in method was performed as previously described^42^. In brief, the crRNA, tracrRNA (#1072532) and Cas9 protein (#1081060) were all purchased from IDT, Coralville, Iowa, USA. The crRNA sequence for PCYT-1::3xFLAG is ATATGAAGTTAATACTCCAA and for GFP::SBP-1 is CAGAATGAACGAAGAATTCG.

The crRNA and tracrRNA were dissolved in IDTE (10mM Tris, 0.1mM EFTA) and IDT- duplex buffer (30mM HEPE, pH7.5; 100mM potassium acetate), respectively, at a concentration of 0.4ug/ul. Regarding to the insertion template, single-stranded DNA oligos were used for PCYT-1::3xFLAG. The sequence was *C. elegans-*codon optimized and ordered from IDT (standard desalting; 4 nmol Ultramer) as the following: GAATCAAAAAAATACGTTTTTAGGTCAAGAGCTGAAAAATATGAAGTTACTTATCATCGTCATCCTTATAGTCGATATCGTGGTCTTTATAGTCACCATCGTGATCTTTGTAGTC ATACTCCAAAGGAGTCTTTGCCTTATTTCTAGAGGACCTTTTCTTCACG. For GFP::SBP-1, an asymmetric-hybrid dsDNA cocktail was generated using pYW1808 as the template and amplified with two sets of primers : sbp-1_repair_F / sbp-1_repair_R and 97.75_gfp_F / 95.75_gfp_R.

### Quantification and statistical analysis

All statistical analyses were accomplished using GraphPad Prism software (RRID:SCR_002798; version 6.0c, La Jolla, CA). The Student’s t-test was performed for statistical analysis and data were considered significantly different when p < 0.05. All representative results are shown as mean ± SEM for at least three independent replicates of experiments.

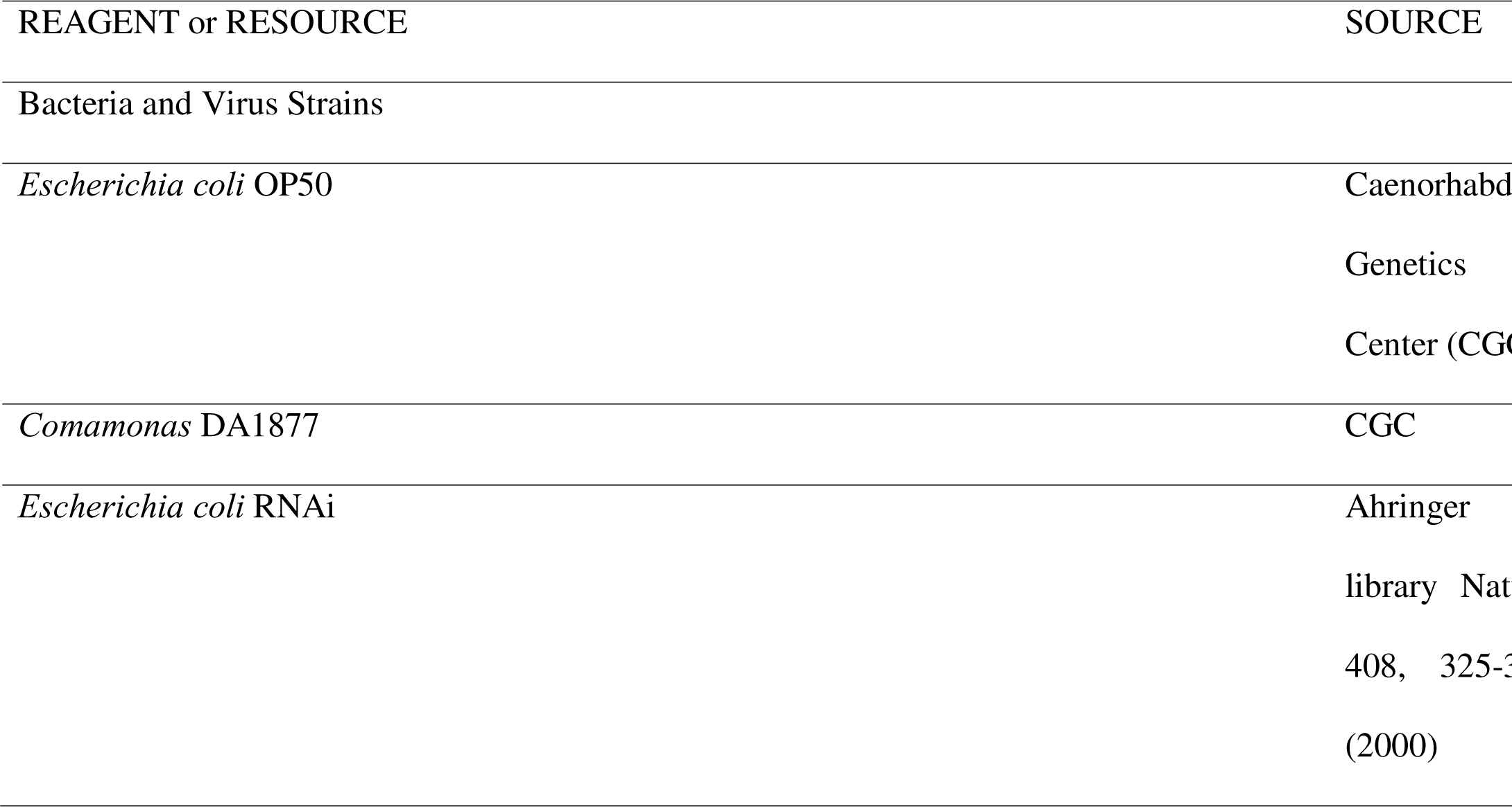
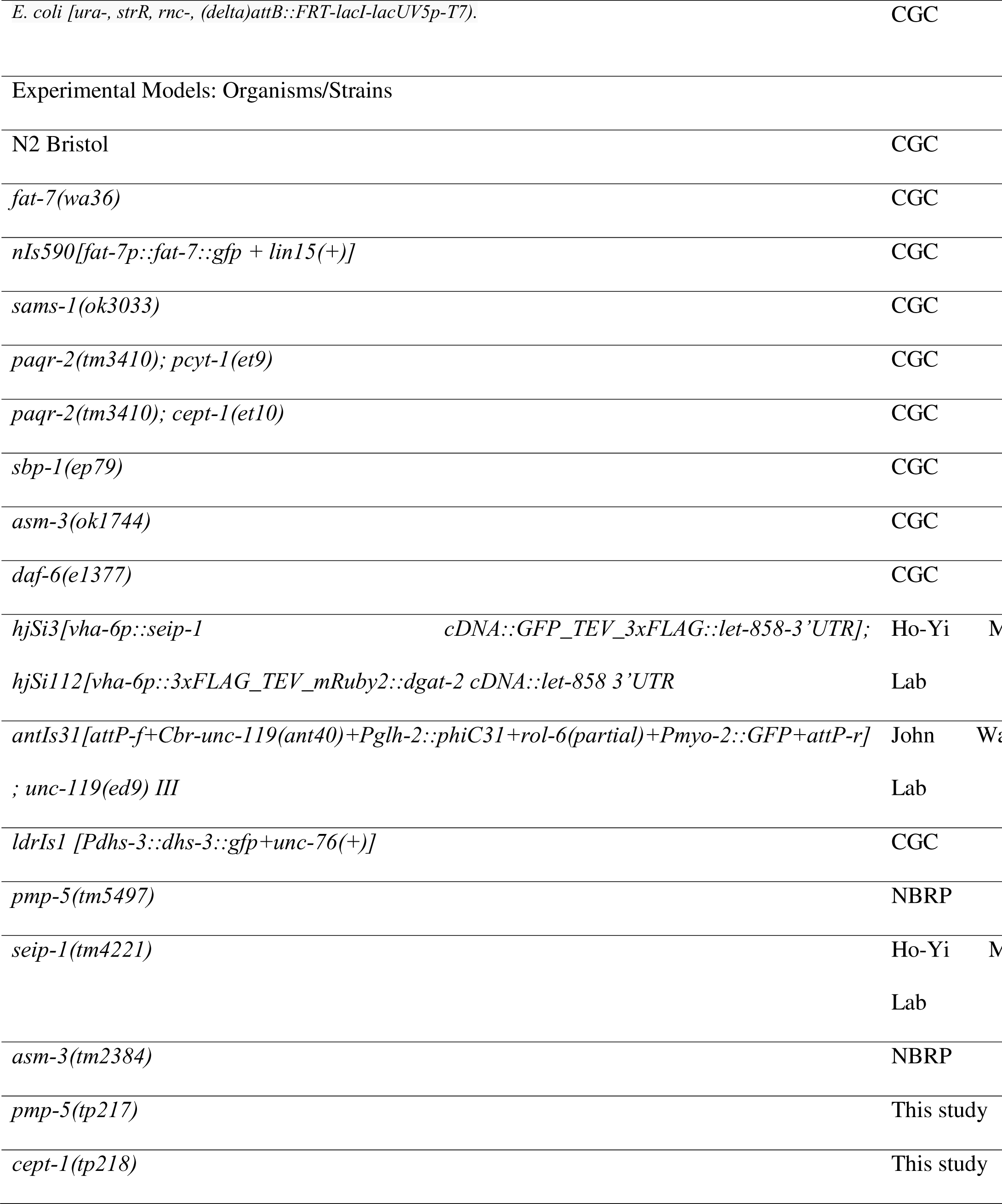
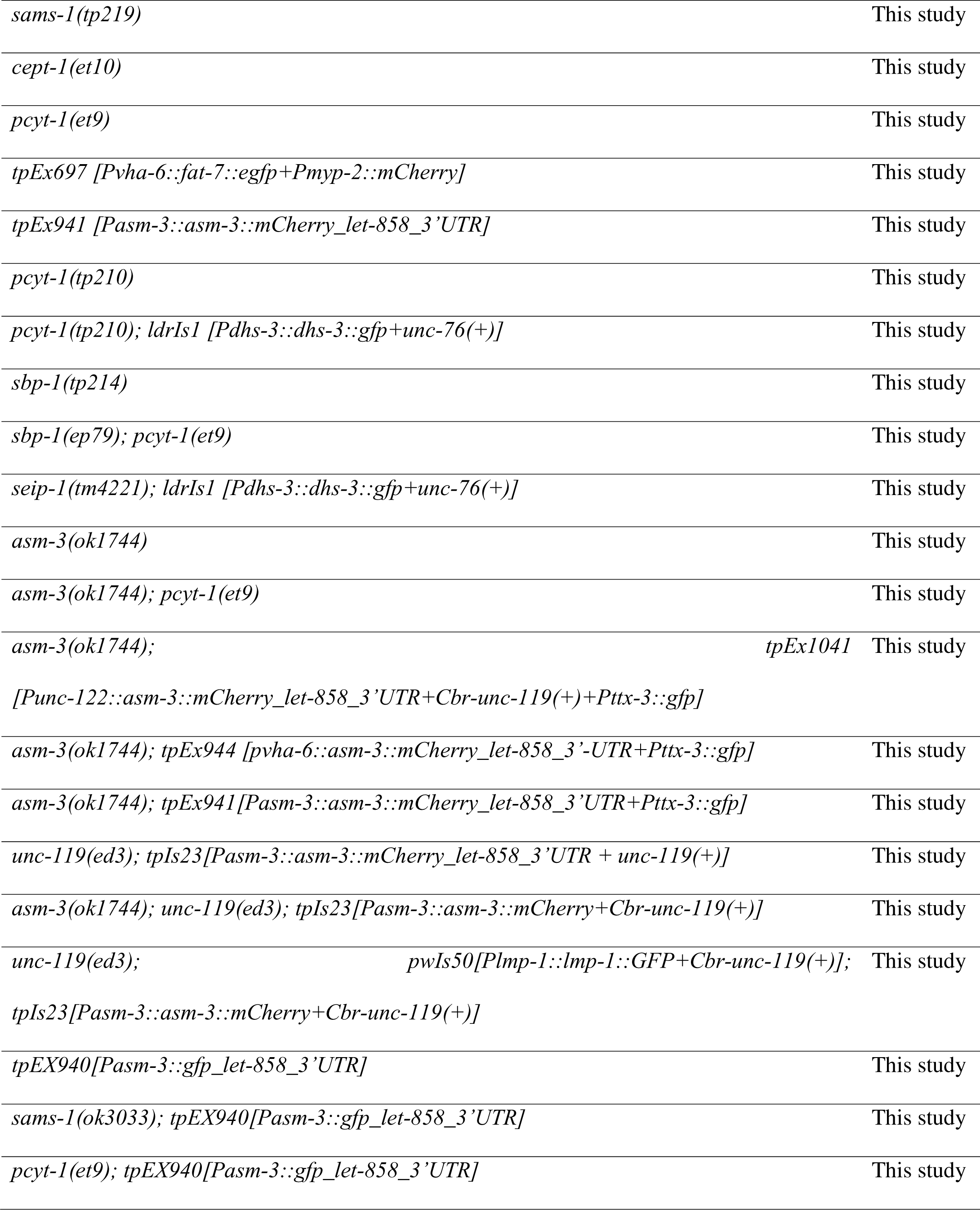
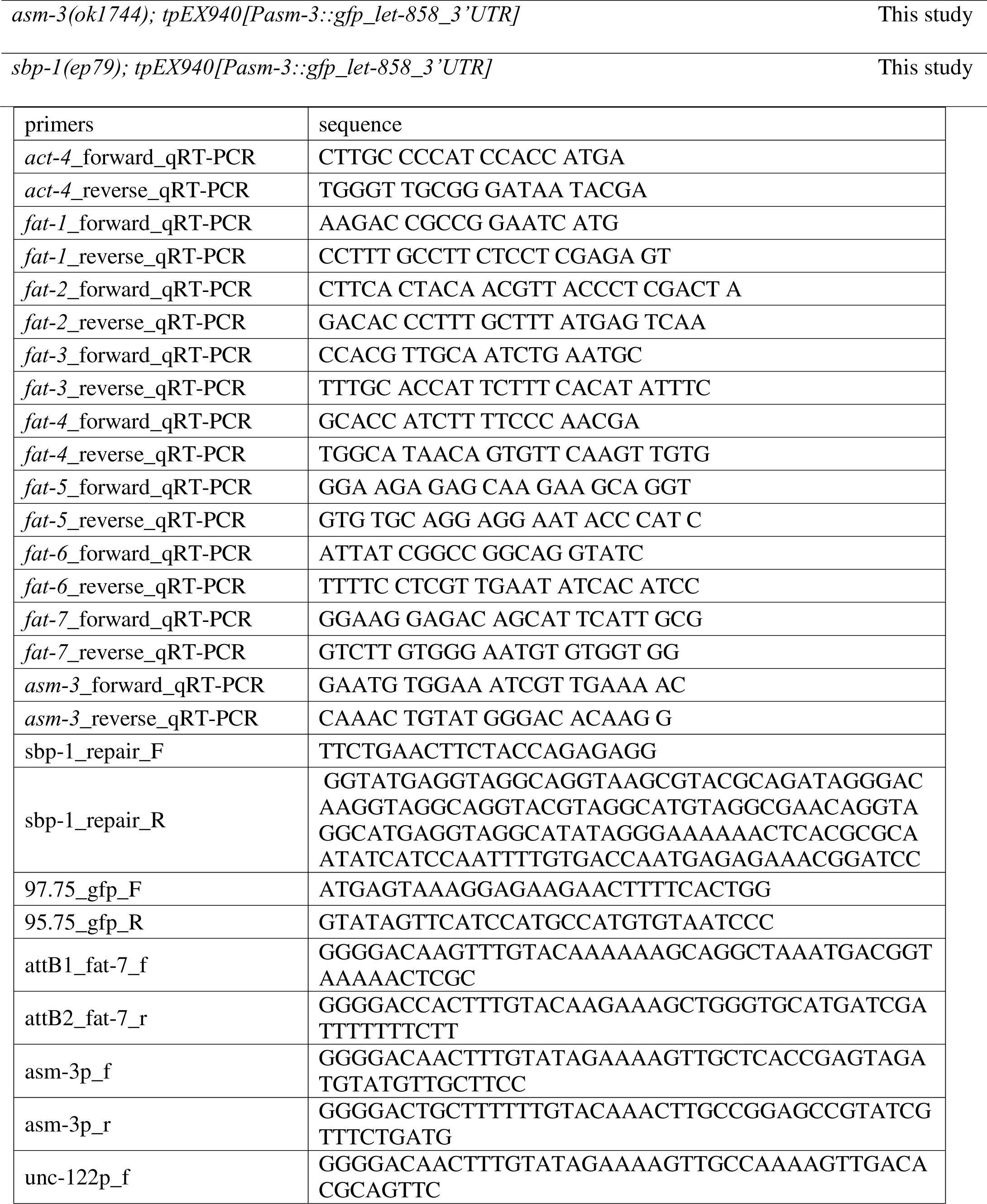
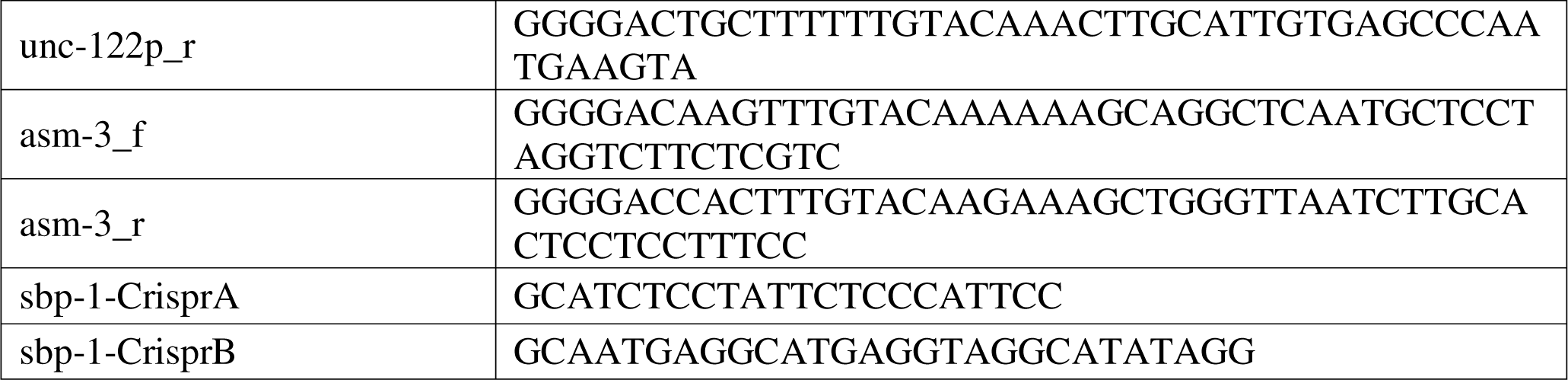

## Acknowledgements

We thank Dr. Ho-Yi Mak of the Hong Kong University of Science and Technology for *C. elegans* strain VS488 for visualization of SEIP-1 and LDs, Dr John Wang at Academic Sinica and Dr. Shih-Peng Chan at National Taiwan University for providing technical supports and materials of the phiC31 recombination system. We also thank the *Caenorhabditis* Genetics Center, supported by a grant from the National Institutes of Health and the National BioResource Project, which is funded by the Japanese government, for providing strains. We thank Technology Commons in College of Life Science and Center for Systems Biology at National Taiwan University, and the *C. elegans* core facility Taiwan, supported by a grant from National Science and Technology Council (NSTC) 111-2740-B-002-002-, for technical and materials supports. We thank Mei-Jane Fang, and Ji-Ying Huang at the Cell Biology Core lab at Institute of Plant and Microbial Biology (IPMB) Academia Sinica for advice on microscopy. The plasmids pML25, pGH112, pADA126, pCR110, and pCFJ159 are kind gifts from Dr. Erik Jorgensen at University of Utah. This work is supported by the Ministry of Science and Technology (Taiwan) (MOST) 110-2311-B-002-009-MY3, NSTC 111-2311-B-002-017- and NSTC 112-2311-B-002-006-grants to Yi-Chun Wu.

