## Supplemental Figures for "Dietary bacteria control *C. elegans* fat content through pathways converging at phosphatidylcholine"

Supplementary data 1

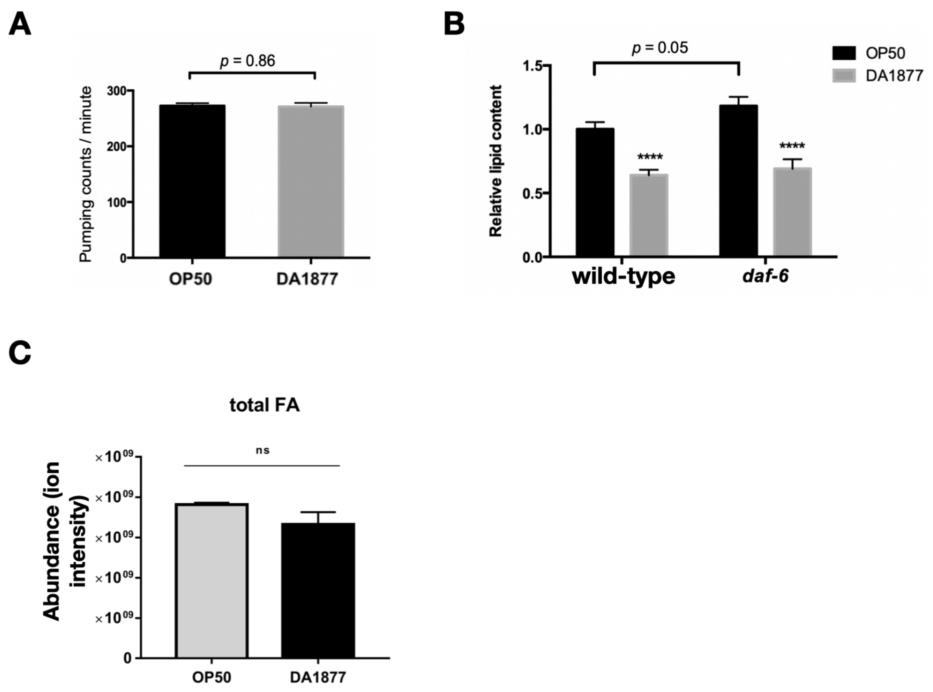

**Supplementary Figure S1. Lipid reduction in DA1877-fed worms is not caused from food intake defect or sensory perception difference**

(A) Pharyngeal pumping rate of wild-type worms fed on OP or DA diet. (B) O.R.O staining results of wild-type and *daf-6(e1377)* worms feeding OP or DA. (C) GC-MS analysis of total fatty acid amount in OP- or DA-fed worms. Statistical analysis was performed using Two-tailed T tests and the data is presented as mean ± SEM. * indicates comparison of dietary effects between OP- and DA-fed worms with the same genotype. $^{****/\#\#\#\#}$ $P < 0.0001$.

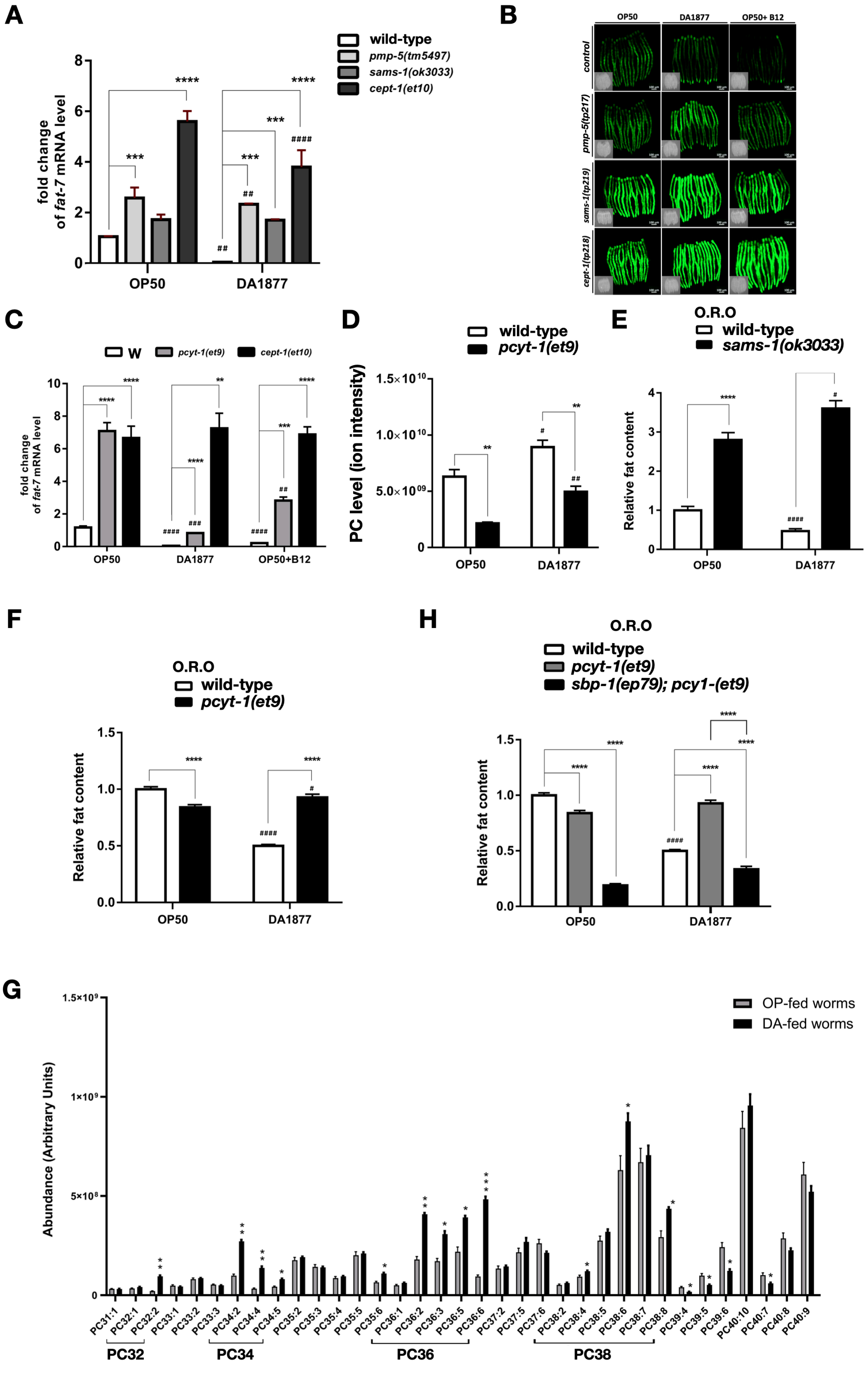
A
wild-type
pmp-5(tm5497)
sams-1(ok3033)
cept-1(et10)
fold change of fat-7 mRNA level
OP50
DA1877
B
OP50
DA1877
OP50+ B12
control
pmp-5(tp217)
sams-1(tp219)
cept-1(tp218)
C
W
pcyt-1(et9)
cept-1(et10)
fold change of fat-7 mRNA level
OP50
DA1877
OP50+B12
D
wild-type
pcyt-1(et9)
PC level (ion intensity)
OP50
DA1877
E
O.R.O
wild-type
sams-1(ok3033)
Relative fat content
OP50
DA1877
F
O.R.O
wild-type
pcyt-1(et9)
Relative fat content
OP50
DA1877
H
O.R.O
wild-type
pcyt-1(et9)
sbp-1(ep79); pcy1-(et9)
Relative fat content
OP50
DA1877
G
OP-fed worms
DA-fed worms
Abundance (Arbitrary Units)
PC32
PC34
PC36
PC38

**Supplementary Figure S2. B12-SAM-PC axis is critical for the decreased *fat-7* expression and fat content in DA-fed worms**

(A) qRT-PCR results of endogenous *fat-7* mRNA level in mutants requested from CGC feeding OP or DA. Experiment contained 3 biological repeats and n = 10 for each repeat. (B) Images for the quantitative results of FAT-7::GFP signals in the control worms DMS303(*nIs590)* and mutants isolated from screen feeding OP, DA, or OP with B12 supplementation in Figure 2H. (C) qRT-PCR results of *fat-7* mRNA level in wild-type*, pcyt-1(et9),* and *cept-1(et10)* feeding OP, DA or OP supplemented with B12. Experiment contained 3 biological repeats and n = 10 for each repeat. (D) LC-MS analysis showing PC level in wild-type and *pcyt-1(et9)* worms feeding OP or DA. (E, F) O.R.O staining results of wild-type, *sams-1(ok3033)* and *pcyt-1(et9)* worms feeding OP or DA. Experiment contains 3 biological repeats and n ≥ 50 in each repeat. (G) LC-MS analysis showing level of individual PC species in wild-type worms feeding OP or DA. (H) O.R.O staining results of wild-type, *pcyt-1(et9),* and *sbp-1(ep79); pcyt-1(et9)* feeding OP or DA diet. Experiment contains 3 biological repeats and n ≥ 50 for each repeat. Statistical analysis was performed using Two-tailed T tests and the data is presented as mean ± SEM. # indicates comparison of dietary effects between OP- and DA-fed worms with the same genotype. * indicates comparison between different strains feeding the same diet. $^{*/\#}$ $P < 0.05$, $^{**/\#\#}$ $P < 0.01$, $^{***/\#\#\#}$ $P < 0.001$, $^{****/\#\#\#\#}$ $P < 0.0001$.

### Supplementary data 3

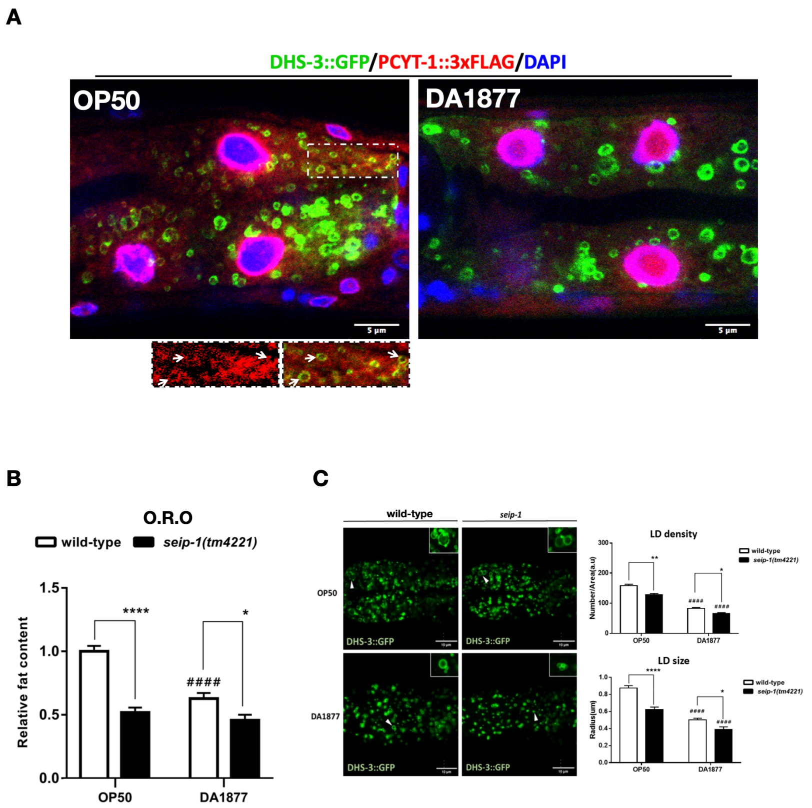

**Supplementary Figure S3. SEIP-1 regulates lipid content and LD size and numbers on both diets**

(A) Representative confocal images of PCYT-1::3xFLAG (Red) in the intestinal cells of worms feeding OP or DA with DAPI staining (Blue) and LD marker, DHS-3::GFP (Green). Scale bar = 5μm. Insets are magnified views from white dotted rectangle. Arrows indicate co-localization of PCYT-1 and LDs. (B) O.R.O staining results of wild-type worms and *seip-1(tm4221)* mutants feeding OP or DA. Experiment contains 3 biological repeats and n ≥ 50 in each repeat. (C) Representative images for visualization of intestinal LDs in wild-type and *seip-1(tm4221)* worms containing the LD marker DHS-3::GFP. Right panel showing quantitative results of numbers and the size of DHS-3::GFP-labeled LDs using MATLAB software. Statistical analysis was performed using Two-tailed T tests and the data is presented as mean ± SEM. # indicates comparison of dietary effects between OP- and DA-fed worms with the same genotype. * indicates comparison between different strains feeding the same diet. $^{*/\#}$ $P < 0.05$, $^{**/\#\#}$ $P < 0.01$, $^{***/\#\#\#}$ $P < 0.001$, $^{****/\#\#\#\#}$ $P < 0.0001$.

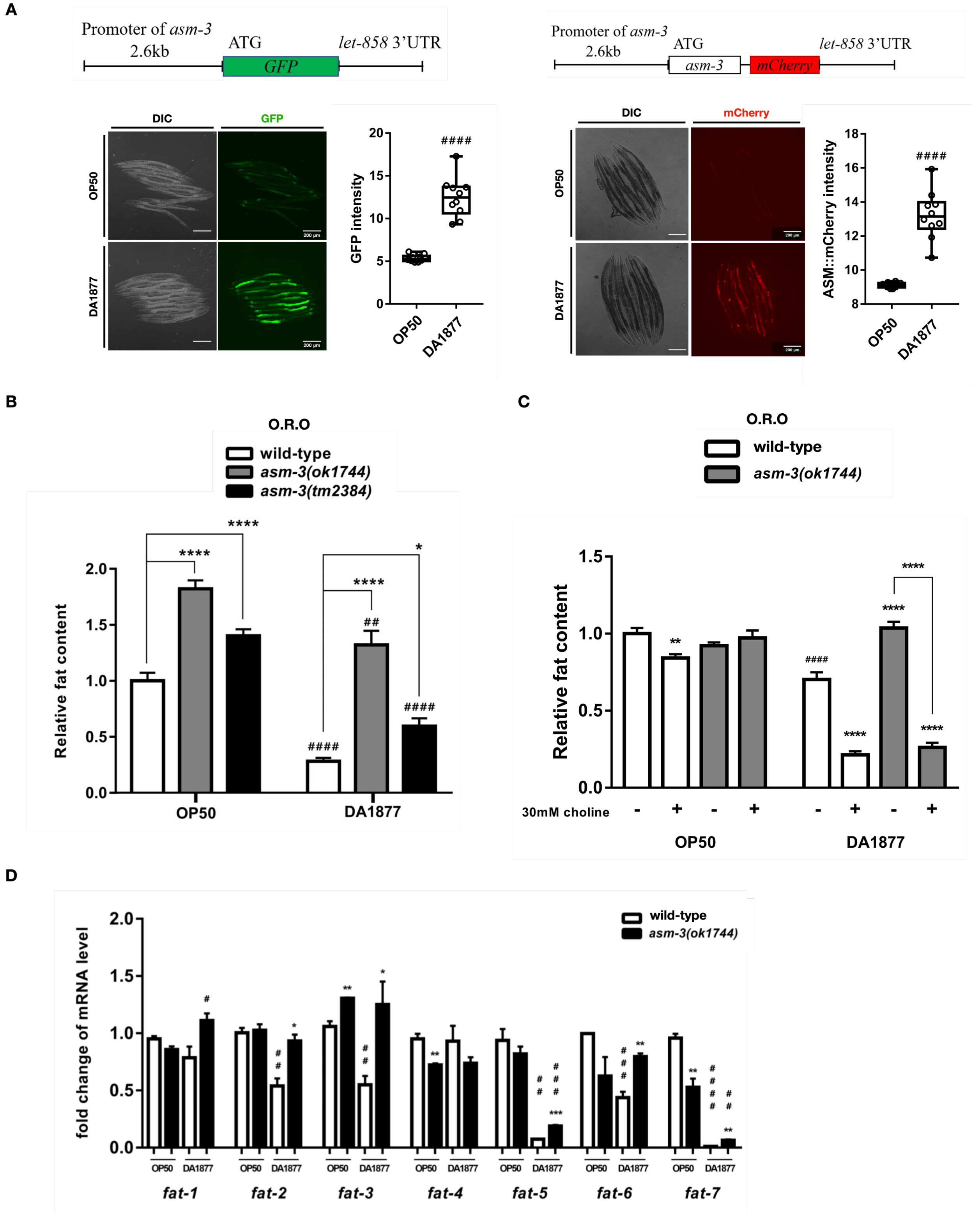
A
Promoter of asm-3
2.6kb
ATG
let-858 3'UTR
GFP
Promoter of asm-3
2.6kb
ATG
asm-3
mCherry
let-858 3'UTR
DIC
GFP
OP50
DA1877
GFP intensity
####
DIC
mCherry
ASM::mCherry intensity
####
B
O.R.O
wild-type
asm-3(ok1744)
asm-3(tm2384)
Relative fat content
****
*
##
####
OP50
DA1877
C
O.R.O
wild-type
asm-3(ok1744)
Relative fat content
30mM choline
OP50
DA1877
D
wild-type
asm-3(ok1744)
fold change of mRNA level
fat-1
fat-2
fat-3
fat-4
fat-5
fat-6
fat-7

**Supplementary Figure S4. The expression of *asm-3* is upregulated in response to DA and loss of *asm-3* promotes the expression of most fatty acid desaturases**

(A) The cartoon in the upper panel showing the design of *asm-3* transcriptional and translational reporters. Images and quantitative results in the lower panel showing the level of GFP and mCherry signal in the AMS-3 transcriptional and translational reporters respectively feeding OP or DA. The centerline in the box denotes the median value. (B) O.R.O staining results of wild-type*, ams-3(ok1744)* and *asm-3(tm2384)* mutants feeding OP or DA. (C) O.R.O staining results of wild-type and *ams-3(ok1744)* mutants feeding OP or DA supplemented with 30 mM choline. (D) qRT-PCR results of mRNA level of *fat-1, 2, 3, 4, 5, 6, ,7* in wild-type and *asm-3(ok1744)* mutants feeding OP or DA. Statistical analysis was performed using Two-tailed T tests and the data is presented as mean ± SEM. # indicates comparison of dietary effects between OP- and DA-fed worms with the same genotype. * indicates comparison between different strains feeding the same diet. $^{*/\#}$ $P < 0.05$, $^{**/\#\#}$ $P < 0.01$, $^{***/\#\#\#}$ $P < 0.001$, $^{****/\#\#\#\#}$ $P < 0.0001$.

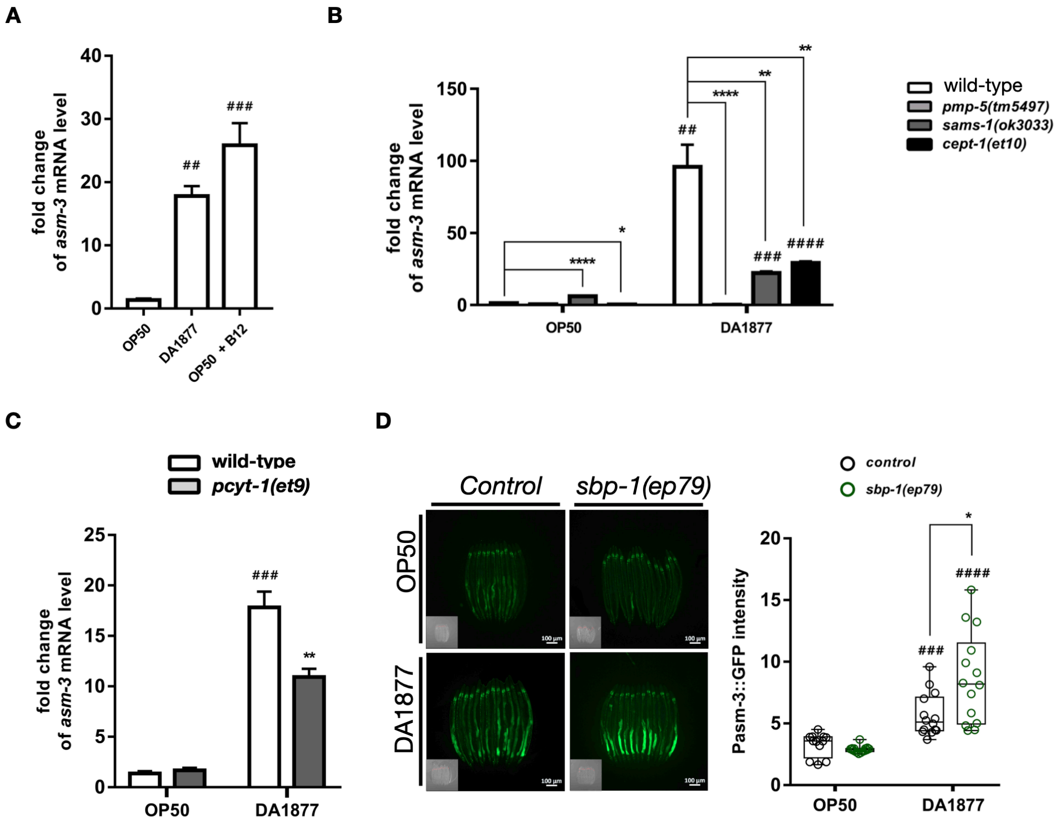
A
B
C
D
fold change of asm-3 mRNA level
OP50
DA1877
OP50 + B12
##
###
wild-type
pmp-5(tm5497)
sams-1(ok3033)
cept-1(et10)
****
**
*
####
pcyt-1(et9)
Control
sbp-1(ep79)
control
Pasm-3::GFP intensity
100 µm

**Supplementary Figure S5. *asm-3* is induced by B12-SAM-PC axis on DA diet**

(A) qRT-PCR results of *asm-3* mRNA level in worms feeding OP, DA or OP supplemented with B12. (B) qRT-PCR of *asm-3* mRNA level in wild-type worms and B12-SAM-PC axis mutants. (C) qRT-PCR of *asm-3* mRNA level in wild-type and *pcyt-1(et9)* worms feeding OP or DA. (A-C) Experiment contained 3 biological repeats and n = 10 for each repeat. (D) Images and quantitative results of the signals of *asm-3* transcriptional reporter in *control* and *sbp-1(ep79)* mutant background. The centerline in the box indicates the median value in the data. Statistical analysis was performed using Two-tailed T tests and the bar plot is presented as mean ± SEM. # indicates comparison of dietary effects between OP- and DA-fed worms with the same genotype. * indicates comparison between different strains feeding the same diet. $^{*/\#}$ $P < 0.05$, $^{**/\#\#}$ $P < 0.01$, $^{***/\#\#\#}$ $P < 0.001$, $^{****/\#\#\#\#}$ $P < 0.0001$.

### Supplementary Figure 6

**A**

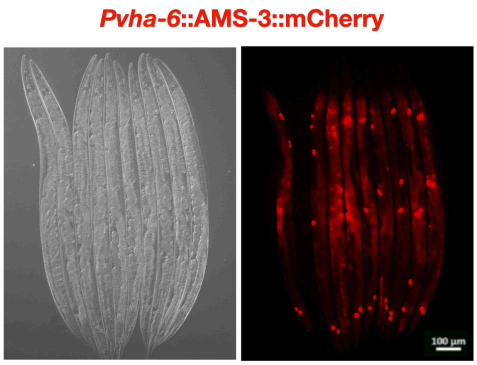


**B**

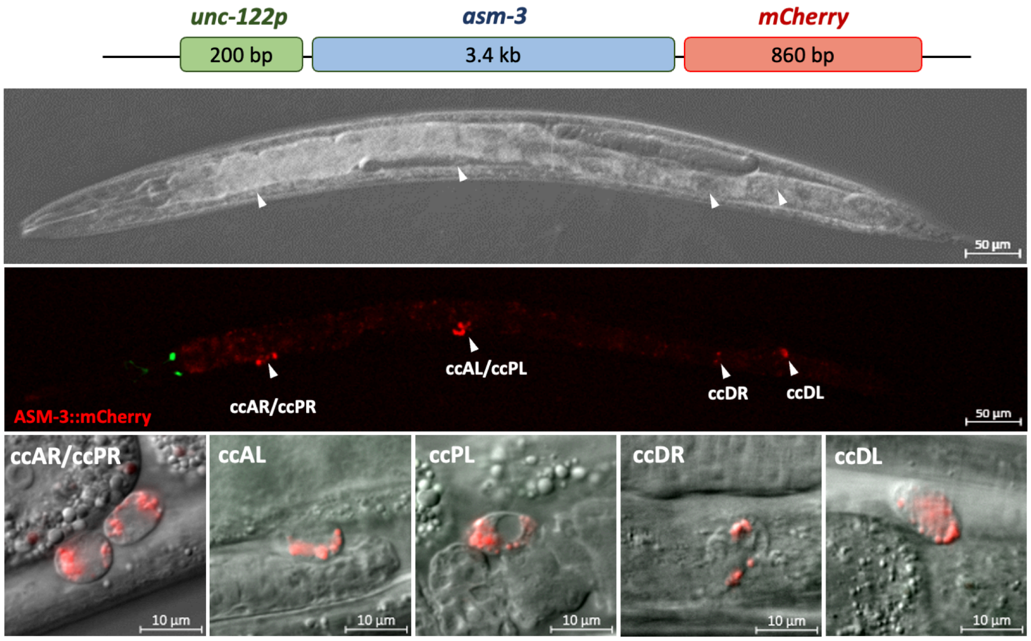

**Supplementary Figure S6. The expression pattern of ASM-3::mCherry driven by intestine-specific promoter and coelomocyte-specific promoter**

(A) Images of ASM-3::mCherry signals driven by intestinal *vha-6* promoter in the worms. (B) The cartoon showing the design of coelomocytes-expressed *asm-3* construct driven by *unc-122* promoter. Images in the lower panel showing localization of ASM-3 in *asm-3(ok1744); tpEx1041* worms expressing *asm-3::mCherry* driven by coelomocyte promotor *unc-122*. White arrowheads indicate coelomocytes.
